# Neuronal activity modulates the incorporation of newly translated PSD-95 into a robust structure as revealed by STED and MINFLUX

**DOI:** 10.1101/2023.10.18.562700

**Authors:** Clara-Marie Gürth, Maria Augusta do Rego Barros Fernandes Lima, Victor Macarrón Palacios, Jasmine Hubrich, Angel Rafael Cereceda Delgado, Nikolaos Mougios, Felipe Opazo, Elisa D’Este

## Abstract

The postsynaptic density component PSD-95 undergoes activity-dependent plasticity mechanisms that rely on protein synthesis and structural remodeling. How synaptic activity can influence these dynamics at the single synapse level remains unclear. Here we combine genome-editing, pulse-chase experiments, STED and 3D MINFLUX nanoscopy on hippocampal neuronal cultures to study the integration of newly translated PSD-95 molecules at postsynaptic sites and their rearrangement within individual clusters at near-molecular resolution. We show that the amount of newly translated PSD-95 recruited to individual synapses scales with synaptic size, and modulates in a bidirectional manner, resulting in less new protein following excitatory and more new protein following inhibitory stimulation. Furthermore, we show that within synaptic clusters PSD-95 has a dispersed organization that is largely robust to long-lasting changes in activity. Altogether, this work sheds new light on the mechanisms underlying plasticity at the single synapse level, adding previously inaccessible information.

## Introduction

Synaptic scaling is a homeostatic plasticity mechanism induced at the level of synaptic sites, the smallest functional units in memory formation, by long-lasting changes in activity. It is supported by new protein synthesis and mediated through changes in postsynaptic AMPA receptor accumulation (*1–3*).

Post Synaptic Density 95 (PSD-95) is a key component of the excitatory postsynapse functioning as a scaffold to establish the positioning of AMPA and NMDA receptors (*4–6*). PSD-95 facilitates synaptic scaling by adapting both the size and the geometry of the scaffold. Changes of the post synaptic density (PSD) size require variation in the protein levels. Indeed, protein degradation and translation bidirectionally modulate PSD-95 protein levels during synaptic scaling (*7–9*), as proved by biochemical methods at the whole-culture level (*9, 10*). Newly translated PSD-95 molecules are incorporated at the postsynapse, and electron microscopy data on a small number of synapses suggest that these molecules do not only replace the old ones but also exhibit a distinct localization and possibly function (*7*). However, this hypothesis has so far not been verified, and a comprehensive, comparative analysis of the accumulation and localization of newly synthesized PSD-95 proteins *in situ* at the single synapse level in the context of synaptic scaling is lacking.

Changes in PSD-95 geometry support plasticity by tuning synaptic transmission (*11–15*). This is achieved by dynamically influencing the alignment of postsynaptic receptors with presynaptic release sites (*16*). At the nanoscale, PSD-95 clusters in 80-180 nm-sized nanodomains, which can be considered as PSD basic structural units or building blocks (*12, 17, 18*). The majority of the synaptic sites exhibit a single nanodomain (*12, 17*). However, the number of these structures is influenced by synaptic scaling induced by a persistent inhibitory (but not excitatory) stimulation (*12, 17*). Recent 3D MINFLUX and expansion microscopy experiments on a limited number of synaptic sites suggest the presence of a quasi-regular PSD-95 organization within the nanodomains (*19, 20*). Nevertheless, this evidence still has to be confirmed on a larger dataset and the impact of synaptic scaling on this organization is unknown.

Here, we dissect the two mechanisms through which PSD-95 reacts to long-lasting inhibitory and excitatory stimulation at sub-synaptic level: (i) we perform pulse-chase experiments on endogenously HaloTag-tagged PSD-95 in cultured neurons followed by quantitative multicolor STED nanoscopy to analyze the amount and distribution of newly synthesized proteins; (ii) we used 3D MINFLUX to analyze the organization of PSD-95 with single-digit isotropic resolution.

## Results

### Newly translated and old PSD-95 do not mix homogeneously at synaptic site

We set out to examine PSD-95 de *novo protein* synthesis at individual synaptic sites preserving both molecular specificity and spatial information. To this end, we first established a pulse-chase strategy using spectrally separated, live-cell compatible HaloTag substrates applied to neuronal cultures at different time points (*21, 22*). In detail, the chimeric PSD-95-HaloTag protein in the cells was labeled to saturation with the first fluorescent Halo substrate (pulse, 580CP-Halo, 1 µM for 2 h). Cultures were then incubated for two days before adding a second HaloTag substrate to label newly translated protein (chase, SiR-Halo, 0.5 µM for 24 h, **Fig. 1a, b**). This system was implemented on CRISPR/Cas9 endogenously tagged rat hippocampal neurons to guarantee the expression of the PSD-95-HaloTag from the native promoter (*23*). The completeness of both the pulse and chase labeling steps was tested by applying the respective other Halo substrate without any intermediate incubation, demonstrating that all available HaloTags were successfully labeled (**Fig. S1a**).

We next evaluated the new protein synthesis of PSD-95 under basal conditions in multi-color STED images of pulse-chase labeled mature neuronal cultures (DIV 15-20). STED images showed signal of both, old and new PSD-95-HaloTag in the majority of synaptic sites and an overlap of the two channels, although some intensity clusters in either the old or new protein channel were distinguishable (**Fig. 1b, c**). To test whether the two protein pools intermix or segregate, we performed a Pearson colocalization analysis. The results of the analysis did not show a clear overlap of the proteins, suggesting that they are not completely colocalizing (**Fig. 1d**). In fact, the area overlap of the two channels was approximately 50%, indicating that the signal of the new protein (SiR) occupied half of the area of the signal of the old protein (580CP, **Fig. S1b**). However, this might be ascribed to the fact that 580CP delivers a lower resolved STED image than SiR due to the smaller cross-section with the STED beam, with consequently larger PSD-95 area in the 580CP channel. To rule out this option, we generated control images in which the SiR channel was artificially deteriorated with a 3-pixel gaussian filter to compare the same signal at different resolutions (control images). In these control images the overlapping area of the two channels was 100% (**Fig. S1b**). We then compared the Pearson correlation coefficients of pixels which have a non-zero value in both channels of real images and observed a correlation of around 50%, compared to 75% in generated control images and ∼80% in control multicolor beads (**Fig. 1e**). Thus, these controls and data suggest that although newly translated PSD-95 proteins are present in the entire PSD, they do not homogenously mix with the old protein pool.

**Fig. 1.**
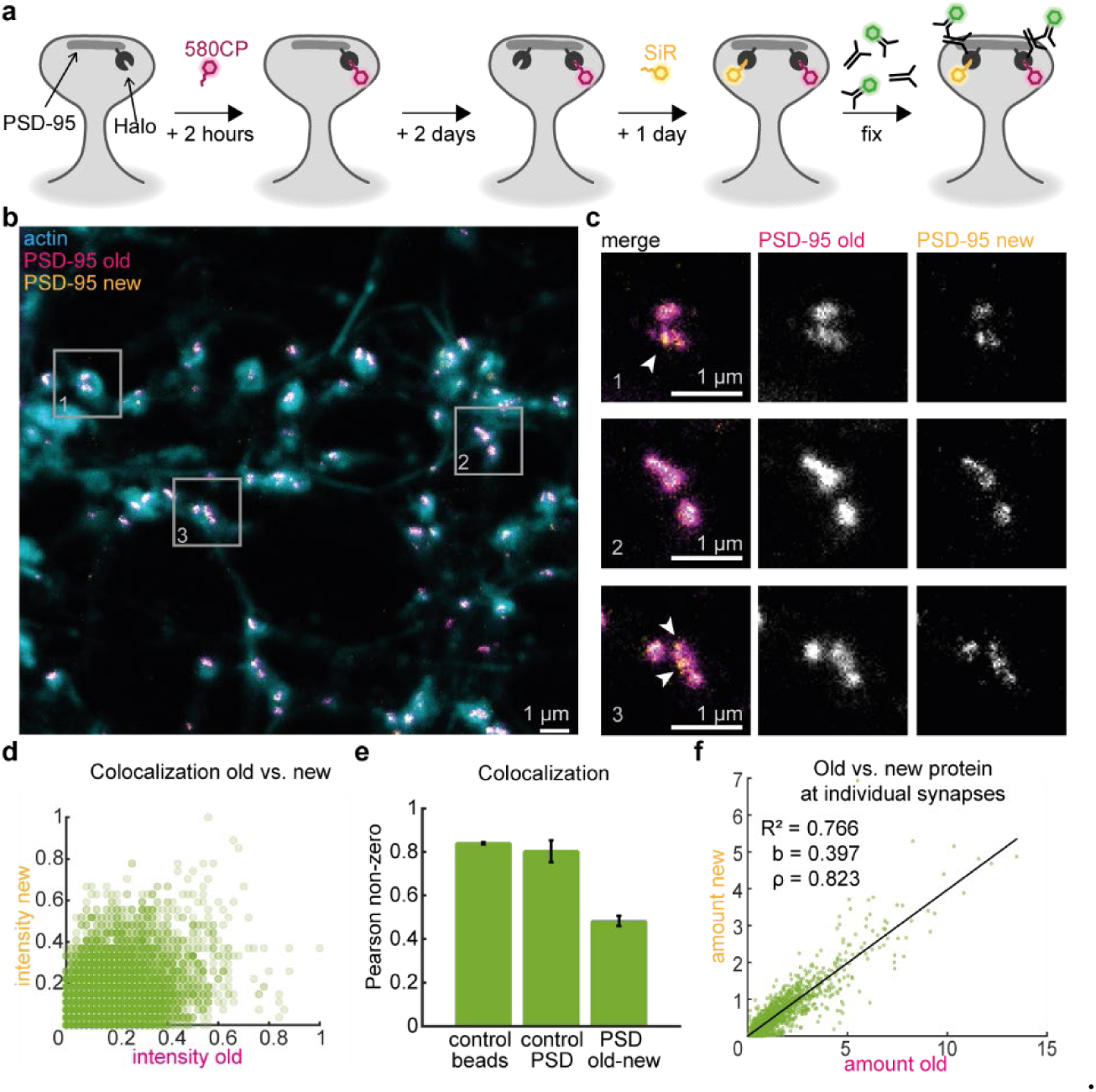
Pulse-chase labeling for *in situ* visualization of endogenous PSD-95 protein translation. (**a**) Experimental workflow of pulse-chase labeling of HaloTag-tagged PSD-95 in neuronal cultures. Primary hippocampal neurons endogenously expressing PSD-95-HaloTag were labeled with a first Halo substrate (pulse, magenta, 580CP) for 2h, then washed and kept in culture for 2 more days. Newly translated PSD-95-HaloTag proteins were then labeled with a second Halo substrate (chase, yellow, SiR) for 24 h, followed by fixation and immunolabeling with an antibody against HaloTag (green) and STED imaging. (**b**) Representative dual color STED image of pulse-chased neuronal cultures. Grey boxes represent synapses shown in (c). (**c**) Close-up of selected synaptic sites. Arrowheads point at regions where an accumulation of the new PSD-95 does not colocalize with the old PSD-95. (**d**) Colocalization of old and new protein in dual color STED images in normalized pixel intensity correlation (Pearson colocalization analysis, calculated for the image shown in b). (**e**) Pearson non-zero coefficient of old and new protein compared to multicolor fluorescent beads and a control image in which the resolution of the SiR channel has been blurred to achieve a resolution comparable to 580CP. Mean and SEM values of 0.80 ± 0.05 and 0.48 ± 0.02 for control PSD (N_images_ = 4) and PSD old-new (N_images_ = 15), respectively. (**f**) Correlation of old versus new protein amounts at individual synaptic sites (n_synaptic sites_ = 2148, N_independent cultures_ = 5). Amount (expressed in A.U.) was calculated as the product of pixel area and mean intensity of each individual synaptic site. Black line is a linear fit with slope (b), coefficient of determination (R^2^), and Spearman’s correlation coefficient (ρ).

### The amount of newly translated PSD-95 scales with synapse size

We questioned whether new PSD-95 protein incorporation occurs at constant rates over the entire synapse population. To this aim, we used an antibody against HaloTag to identify all synapses of genome-edited cells and applied a high throughput image analysis workflow to compare the relative amounts of pre-existing and newly translated PSD-95-HaloTag in more than 2000 synapses. Interestingly, the relative protein amount (measured as the area of the synaptic site multiplied by the mean fluorescence intensity, see methods) of old and new PSD-95 strongly correlated in all imaged synaptic sites, regardless of their size (**Fig. 1f**). This correlation shows that bigger synaptic sites incorporate larger quantities of newly translated PSD-95 and smaller PSD sites possess lower levels of new protein. Hence, new PSD-95 protein synthesis depends on and scales up with the size of the postsynapse.

### The accumulation of new PSD-95 at synaptic sites depends on neuronal activity

Having observed a correlation between synapse size and the amount of newly translated PSD-95 under basal conditions, we asked whether the incorporation of new PSD-95 at the PSD is modulated by neuronal activity. To test this hypothesis, we silenced or enhanced neuronal activity by treating cultures with tetrodotoxin (TTX) or gabazine (GZ) for 24 hours concurrently with the HaloTag chase protocol (**Fig. 2a**). We then compared the relative amounts of old and new PSD-95-HaloTag in the different conditions (**Fig. 2b,c**). Although the levels of old protein remained constant in between treatments, there was a significant increase in new PSD-95 at synaptic sites in TTX treated cultures and a decrease in new protein in GZ treated cultures compared to control vehicle DMSO treated cultures.

Next, we tested whether the correlation between old and new PSD-95 shifts in treated cultures. We observed strong correlations between the amount of old PSD-95 and new PSD-95 in all conditions (**Fig. 2d-f, Fig. S2**). To better compare the different dataset, we grouped the data into 5 bins with the same number of elements (*24, 25*). Binned data confirmed the strong correlation and highlighted different tendencies towards old or new protein, as reflected by the steepness of the linear fit. Indeed, while for TTX the steepness of the fit was increased compared to the control (b = 0.453 and b = 0.339, respectively), it was reduced for GZ treatment (b = 0.190, **Fig. 2d-f**). The ratios between new and old protein at individual synapses therefore were higher in TTX-treated samples and lower in GZ-treated samples. This indicates an overall higher incorporation of new protein upon synaptic scaling induced by prolonged TTX treatment, and a reduced incorporation of new protein upon prolonged GZ stimulation at the single synapse level.

**Fig. 2.**
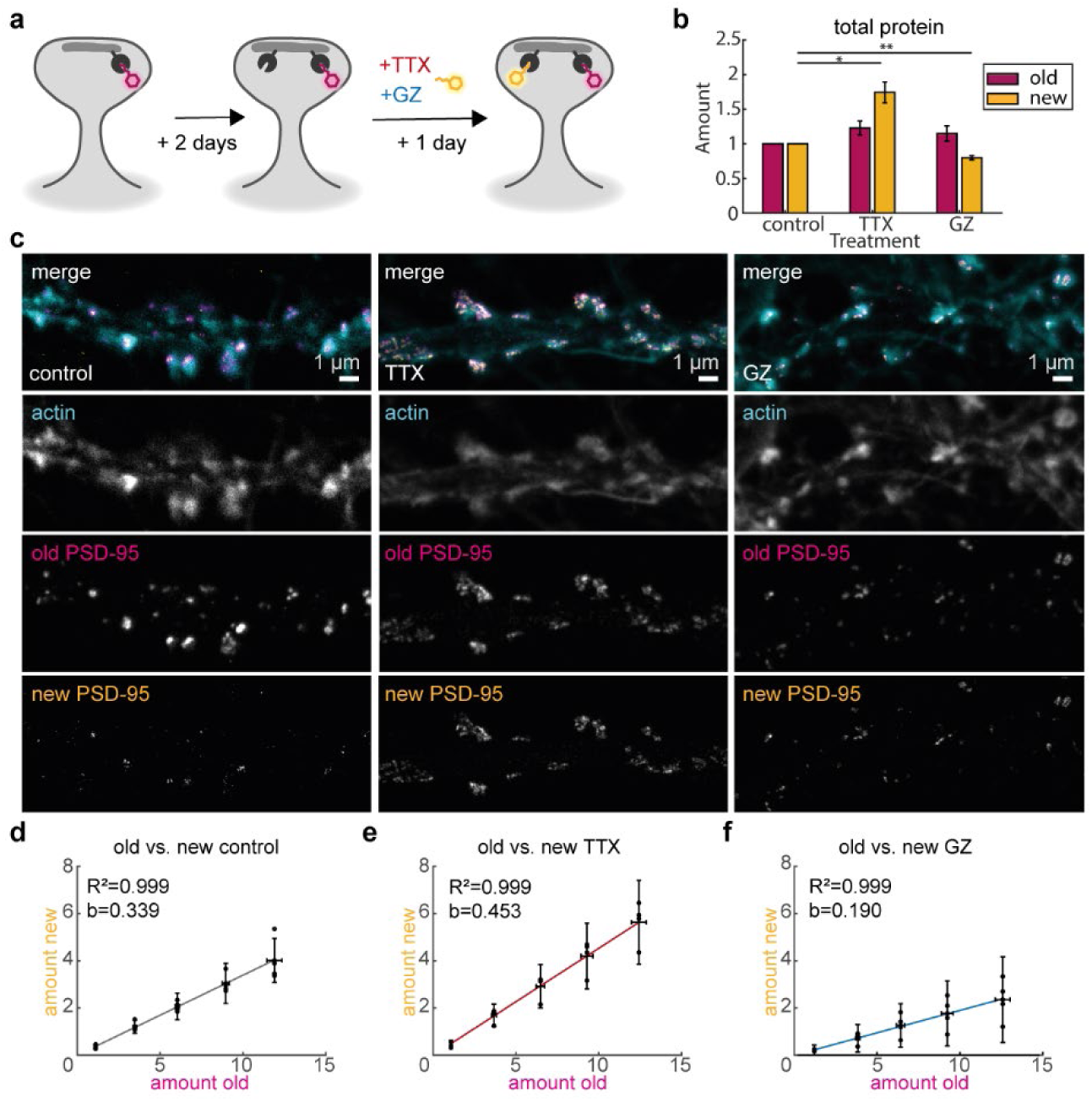
Activity-dependent changes in PSD-95 translation levels. (**a**) Workflow of pulse-chase labeling with prolonged stimulation or silencing of neuronal activity. Primary hippocampal neurons endogenously expressing PSD-95-HaloTag were labeled with the first Halo substrate (pulse, magenta, 580CP), then washed and kept in culture for 2 more days. Newly translated PSD-95-HaloTag protein was then consecutively labeled with a second live cell Halo-substrate dye (chase, SiR, yellow) for 24 hours and, at the same time, cultures were treated with either tetrodotoxin (TTX, 1.5 µM) or gabazine (GZ, 10 µM). (**b**) Quantification of relative overall protein translation. Relative change of old (pulse, magenta) or new (chase, yellow) PSD-95 protein amount (product of pixel area and mean intensity, expressed in A.U.) are plotted as mean of 4 independent experiments, compared to control vehicle (DMSO only). Bars represent the mean and error bars represent SEM values (n_DMSO_ = 1858, n_TTX_ = 1183, n_GZ_ = 790, all from N = 4), where: TTX_old_ = 1.23 ± 0.10, p = 0.22, TTX_new_ = 1.74± 0.15, p = 0.03 and GZ_old_ = 1.15 ± 0.11, p = 0.47, GZ_new_ = 0.80± 0.03, p = 0.01. Statistical analysis was performed with ANOVA followed by Dunnet multiple comparison correction. (**c**) Representative multi-color STED images of dendrites endogenously expressing PSD-95-HaloTag in treated cultures with corresponding actin signal (confocal). Samples were pulse and chased as described in (a) and labeled with phalloidin (actin, confocal) after fixation. (**d, e, f**) Correlation of the amount of old PSD-95-HaloTag protein and new PSD-95-HaloTag protein binned in 5 bins of equal size for control-(d), TTX-(e) and GZ-(f) treated cultures (same dataset as b). Amounts expressed in A.U. Black, red or blue line is a linear fit with slope b and coefficient of determination R^2^. Vertical and lateral error bars indicate SE. Corresponding not-binned data is reported in Fig. S2.

### Highest labeling density for 3D MINFLUX imaging of PSD-95 using nanobodies and DNA-PAINT

Synaptic scaling does not impact only protein turnover but also the overall architecture of PSD-95 (*6, 26*). Therefore, we set out to investigate the nanoscale organization of PSD-95 as a function of activity by using 3D MINFLUX, a technique which provides exquisite localization precision (*27*). First of all, a suitable sample preparation procedure had to be established. To maximize the degree of labeling and minimize the linkage error, we used a nanobody specific for PSD-95 (*28*). Because achieving the single molecule regime required by MINFLUX is challenging and prone to energy transfer in a crowded compartment such as synapses (*29*), we explored different blinking mechanisms. In particular, we compared the number of molecules detected with the anti-PSD-95 nanobody conjugated with either two sulfoCy5 fluorophores for conventional photochemical blinking or with a single-stranded DNA for DNA-PAINT MINFLUX imaging (*30*). Identification of the postsynaptic sites was facilitated by phalloidin counterstaining of dendritic spines, and further supplemented by labeling of presynaptic compartments (bassoon) in the DNA-PAINT samples. After filtering of spurious events (see methods for details), the obtained localization precision using 150 photons per localization was of σ_x_= 5.5 ± 0.8 nm, σ_y_= 5.6 ± 0.8 nm, and σ_z_= 2.9 ± 1.2 nm for sulfoCy5 samples, and σ_x_= 5.1 ± 0.5 nm, σ_y_= 5.4 ± 0.5 nm, and σ_z_= 4.4 ± 0.5 nm for DNA-PAINT samples. The localization precision improved to ∼3-3.8 nm when using 500 photons per localization in the DNA-PAINT samples (**Figure S3**). The resolution obtained was therefore significantly higher than that obtained in previous studies analyzing changes in PSD-95 geometry in 2 dimensions (*17, 18*).

To quantify the differences between imaging with sulfoCy5 and DNA-PAINT, individual synaptic sites were identified by DBSCAN (search radius eps = 200 nm, minimum number of points per cluster *minPoints* = 40, **Table S1**, see methods). Using DNA-PAINT, 3 times more synaptic sites per image were detected in comparison to sulfoCy5. Furthermore, the number of individually localized molecules (*i.e.* number of trace identifyer (TID) or number of bursts of localizations stemming from an individual blinking event, **Fig. 3d**) per site was 3 times higher for DNA-PAINT. However, we reasoned that the difference could be due to the repeated localization of the same molecules in DNA-PAINT samples, since closely located individual molecules (*i.e.* TIDs closer than 6 nm), were frequently visualized. These TID could correspond to the repeated localization of the same molecule or to the presence of very closely packed PSD-95 molecules. Since we could not discriminate between these options, to avoid overcounting, all TIDs distancing up to 8.5 nm from each other were grouped and assigned to what we defined as an individual PSD “unit” (**Fig. 3d, Fig. S4**) (*31*). This value corresponds to the Euclidian distance between two points in a 3D space, separated by ∼2σ in each dimension, *i.e.* 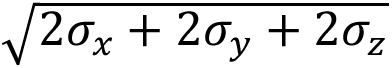. In a conservative approach, the localization precision obtained with 150 photons was considered.

**Fig. 3.**
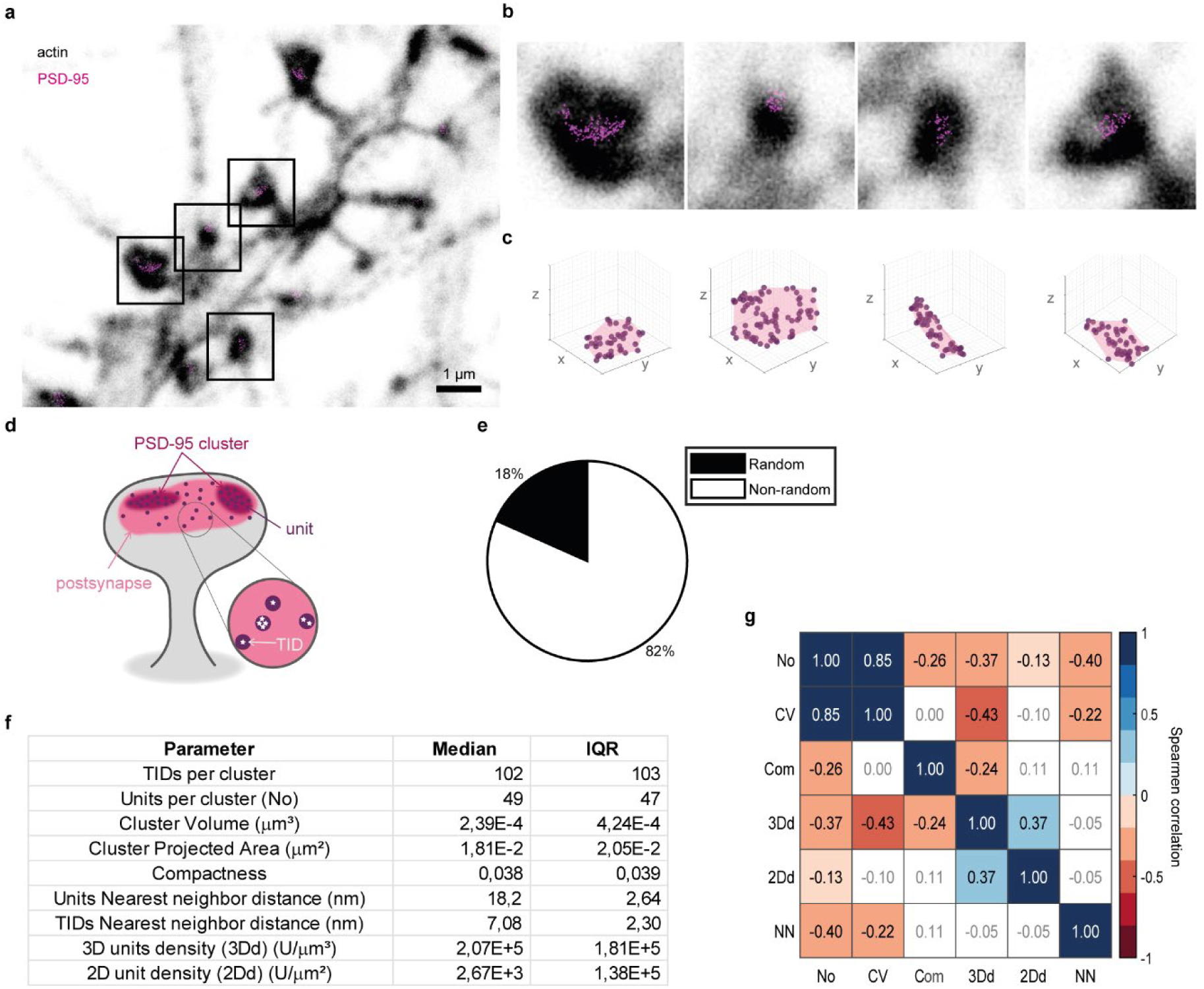
MINFLUX nanoscopy of PSD-95 organization in neuronal cultures under basal conditions. (**a**) Representative image of a dendrite labeled with phalloidin (grey, confocal) and nanobody against PSD-95 (magenta, DNA-PAINT 3D MINFLUX). Grey squares indicate regions shown in b. (**b**) Enlarged images of individual PSD-95 clusters. (**c**) Rendering (light pink) of the alpha shape of the synaptic sites depicted in b. TIDs are shown as gaussian with FWHM of 12 nm, size of each axis is 300 nm. (**d**) Scheme of organizational levels and terminology of PSD-95 considered in this study. A postsynaptic site can have multiple PSD-95 cluster, as determined by a DBSCAN with eps = 75 nm. PSD-95 clusters are formed by PSD-95 units that can contain a single or multiple TIDs (Trace IDs, or individual localizations). (**e**) Percentage of PSD-95 clusters exhibiting a random or non-random (dispersed) spatial organization of PSD-95 units based on the G-function compared to Monte-Carlo simulations. (**f**) Table of quantified parameters for 278 PSD-95 clusters in 3D MINFLUX measurements from 8 cells and 3 independent experimental rounds. (**g**) Correlation matrix of measured parameters using Spearman correlation. Parameters: No – number of TIDs, CV – cluster volume, Com – compactness, 3Dd – unit density in 3D, 2Dd – unit density in 2D projected clusters, NN – units nearest neighbor distance. Blue to red color gradient of the square indicates a positive to negative correlation, as indicated by the number. White squares indicate that the correlation is not significant (threshold 0.05, corresponding p-values in Table S2).

The number of units per synaptic site for DNA-PAINT outperformed sulfoCy5 labeling also in this case. Importantly, with DNA-PAINT neither the number of TIDs per unit nor the number of units increased linearly with the imaging time (minimum imaging time 3 hours), suggesting that most of the nanobodies were detected (**Fig. S5**). Moreover, control samples in which the nanobody was omitted had fewer localizations (no synaptic sites could be detected, data not shown), and the waiting times for a successful localization was 16 times longer than in corresponding samples, indicating a low background due to the free-floating imager strand (**Fig. S6**). Lastly, we compared the efficiency of the PSD-95 nanobody versus a knock-out validated primary antibody, detected using secondary nanobodies and DNA-PAINT (*32*). Also, in this case, the primary PSD-95 nanobody outperformed the antibody regarding TIDs and units per synaptic site (**Table S1**). Therefore, we conclude that using the nanobody in combination with DNA-PAINT provides the best labeling and detection of PSD-95 under our experimental settings using 3D MINFLUX.

### Characterization of PSD-95 with MINFLUX reveals higher-order organization

Having set up a robust way to image PSD-95 at unprecedented resolution, we characterized its organization in cultured neurons under basal conditions. Thanks to the higher quality of the DNA-PAINT nanobody data, more stringent DBSCAN parameters could be applied (eps = 75 nm) to exclusively analyze the PSD-95 molecules located at the PSD, while perisynaptic molecules (*i.e.* adjacent molecules or molecules not lying on the flat postsynaptic plane) were largely excluded (**Fig. S4)**. From here on we will use the term “PSD-95 clusters” to refer to the subset of PSD-95 localizations at synaptic sites which are selected by the more stringent DBSCAN and utilized for the MINFLUX data analysis (**Fig. 3d**). In 3D MINFLUX, PSD-95 clusters showed nearly flat or concave geometries (**Fig. 3a-c, Movie S1**). We obtained a median value of 102 TIDs and 49 units per PSD-95 cluster, with approximately 40% of the units containing more than one TID (**Fig. 3f, Fig. S5b).** Having access to 3D information, enabled us to precisely analyze the volume (median ∼2.4 x 10^-4^ µm^3^) and shape of the PSD-95 clusters (**Fig. 3f**), whose median compactness (∼0.038) indicates a flat geometry. The density of units in the PSD-95 volume was nearly ∼2.1 x 10^5^ U/µm^3^. When looking at the distances between the TIDs, the median value between nearest neighbor distances was 7.1 nm, with the most frequent values for the 1^st^, 2^nd^, and 3^rd^ nearest neighbors being between 2-3 nm for the 1^st^ and 5-6 nm for the 2^nd^ and 3^rd^ nearest neighbors (**Fig. 3f**, **Fig. S7a**). These TIDs could either stem from the same molecule or belong to adjacent molecules, thus considered one unit. At the unit level, the median distance between nearest neighbors in a PSD-95 cluster was 18.2 nm, with the most frequent values peaking at 11-12 nm (**Fig. S7b**). Interestingly, the distribution of the nearest neighbor distances differs from what is expected in the case of spatial randomness (Gaussian-like distribution) and prompted us to assess the presence of an ordered status of PSD-95 molecules with a spatial analysis. To this aim, we applied the Diggle-Cressie-Loosmore-Ford (DCLF) test (*33*) to test for spatial randomness for the units within the confined PSD volume (**Fig. S4**, methods). The results showed with a confidence level α = 0.002 that in 82% of the PSD-95 clusters the units were not randomly distributed, and exhibited a dispersed behavior (**Fig. 3e**). When performing the same analysis at the TID level, 54% of the PSD-95 clusters exhibited a dispersed behavior. This data endorses the hypothesis of a higher-order organization of PSD-95.

Lastly, we wondered whether the analyzed parameters depend on the size of the PSD-95 cluster. To this aim, we tested the parameters for the presence of linear correlations (**Fig. 3g**, **Table S2**). We observed a strong positive correlation between the volume of the PSD-95 cluster and the number of units, indicating that smaller PSDs also have fewer PSD-95 molecules, while larger PSDs have more units. All other parameters showed weak correlations to PSD-95 cluster volume (CV), with the exception of a moderate negative correlation between the 3D density of units and the volume of the cluster (*i.e.* larger clusters have a lower density). This trend is reflected in weak to moderate negative correlations between the number of units per cluster and both the 3D density and nearest neighbor distance of units. Intriguingly, no correlation was observed between the nearest neighbor distance of the units and their 3D density, suggesting that the prevailing distance between units of 11-12 nm is a robust parameter across PSD-95 clusters. Overall, these findings suggest a dispersed organization of PSD-95 at the level of the PSD. This organization is weakly dependent on the size of the postsynapse.

### Detecting nanometer-sized postsynaptic changes

To test the sensitivity of our experimental approach to structural changes in the PSD-95 clusters, we applied 1,6-Hexanediol (Hex) to cultures. Hex is an alcohol widely used to dissolve liquid-liquid phase separation assemblies (*34*) and prolonged (> 10 min) applications to neuronal cultures disrupt synaptic structure (*35*). We opted for a short (2 min) application of 10% Hex, which was sufficient to largely relocalize homer, another PSD scaffolding protein, to dendrites but has minimal impact on PSD-95 (**Fig. S8a-c**). Upon Hex treatment, synaptic sites showed a larger and rounder structure, compatible with the initial phases of a dissociation of molecules from the postsynaptic density (**Fig. S8**). Albeit more units with a comparable number of TIDs and with a shorter median nearest neighbor distance were observed, this effect, undetectable by 2D imaging, was associated with a decrease in the overall 3D density of units per cluster. This data, together with the evidence of a slightly reduced non-randomness in Hex-treated samples (**Fig. S8**), suggests that discrete groups of molecules may dissociate from the PSD during the initial stages of Hex treatment. Taken together, these experiments demonstrate the sensitivity of MINFLUX to detect nanometer-scale postsynaptic changes.

### PSD-95 3D organization is robust to activity changes

Having characterized the nanoscale architecture of PSD-95 clusters in basal conditions and proved that fine changes in their structures can be detected with MINFLUX, we investigated whether prolonged synaptic activity modifies the PSD-95 nanostructure. To this aim, we performed 3D MINFLUX measurements on cultures treated with DMSO (control), TTX or GZ for 24 hours (**Fig. 4**). Under all conditions, 3D MINFLUX imaging revealed PSD-95 clusters. Furthermore, the descriptors of PSD-95 arrangement in DMSO treated samples were comparable to the values obtained for untreated cultures (compare **Fig. 3f** to **Fig. 4**, **Fig. S9**, and **Fig. S10**), proving the robustness of the data. Under all conditions, PSD clusters had comparable median numbers of units and of TIDs per clusters, and a similar volume, although clusters of TTX-treated samples were flatter (compactness DMSO = 0.05, TTX = 0.04, GZ = 0.05, **Fig. 4g**, **Fig. S9)**. At the level of the individual TIDs, TTX-treated samples exhibited a lower 3D density, which was associated with a 1 nm larger distance between the individual TIDs, even if the number of TID per unit was comparable and the distribution of all nearest neighbor TIDs did not differ (**Fig. S10 and Fig. S11**). However, this latter difference was not reflected at the unit level. Indeed, for all conditions the 3D density and median distance between the units (DMSO = 17.6 nm, TTX = 18.0 nm, GZ = 17.8 nm) remained constant (**Fig. 4g**, **Fig. S11**). When looking at the correlation between the analyzed parameters, no major changes were observed between control and treated samples (**Fig. S12**). Importantly, a higher fraction of PSD-95 clusters exhibited a dispersed, non-random distribution in TTX-treated samples (**Fig. 4**h), suggesting the presence of nanoscale rearrangements happening in the postsynaptic densities following activity inhibition. All in all, the nanoscale organization of the PSD-95 clusters appears as largely robust to long-lasting excitatory and inhibitory activity changes.

**Fig. 4.**
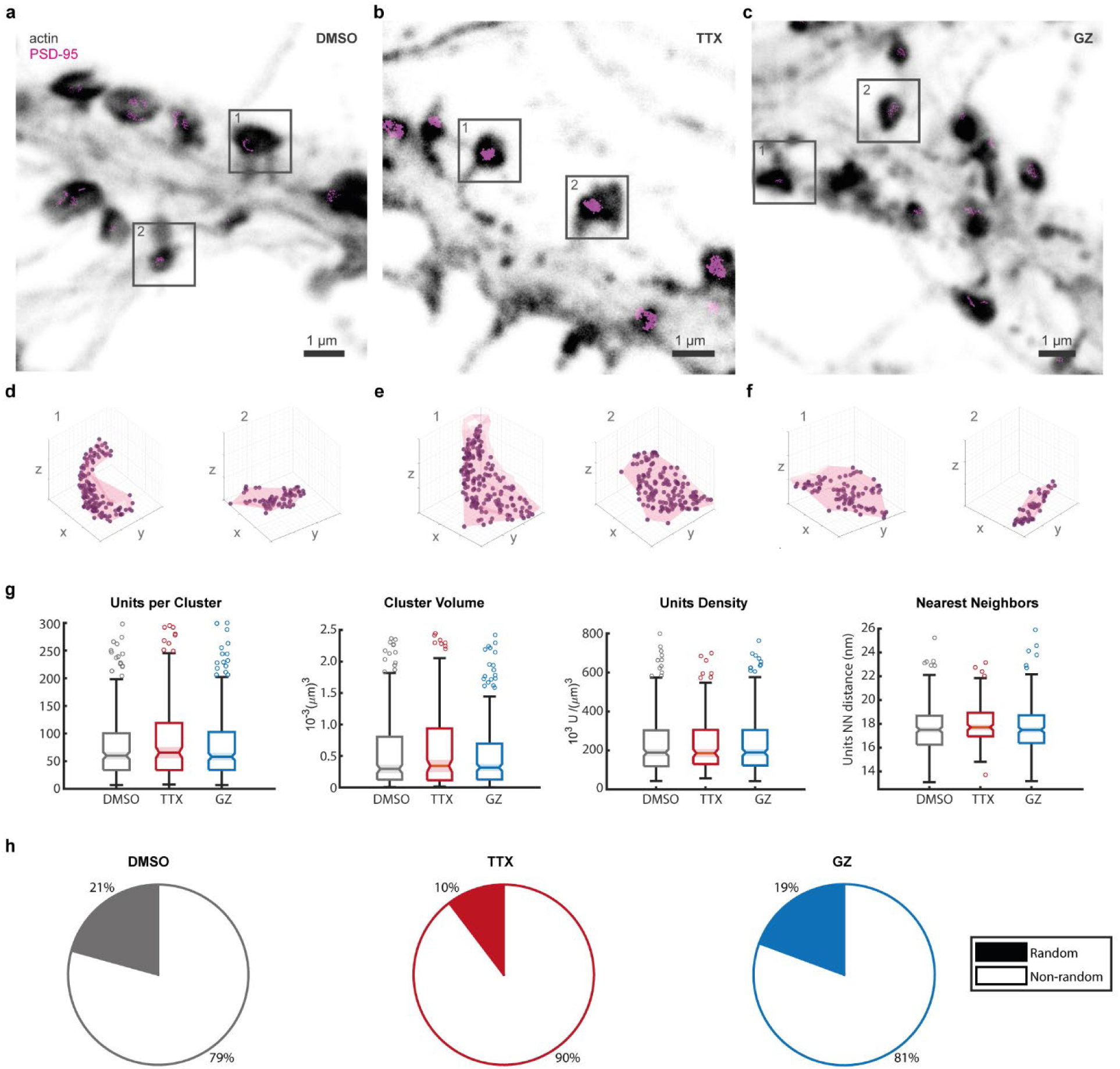
MINFLUX analysis of activity-dependent changes in PSD-95 organization. (**a**) Representative images of dendrites labeled with phalloidin (grey, confocal) and nanobody against PSD-95 (magenta, DNA-PAINT 3D MINFLUX) from cultures treated with (**a**) DMSO, (**b**) TTX or (**c**) GZ for 24 hours prior fixation. Grey squares indicate regions of 3D MINFLUX measurements shown below. (**d-f**) Alpha shape rendering of selected PSD clusters. Axes are 300 nm long. Rendered are TID with an approximate radius of 12 nm. (**g**) Quantification of units per cluster, cluster volume, unit density and units nearest neighbor distances of PSD-95 clusters in 3D MINFLUX measurements with different treatments (numbers of analyzed PSD clusters: DMSO = 261, TTX = 174, GZ = 279 from 8-9 cells and 3-4 independent experimental rounds). Box-plots include the data between the 25^th^ and 75^th^ percentile, median and confidence interval are rendered. Whiskers extend to most extreme data points and outliers are plotted as circles. Statistical analysis performed using Linear Mixed Model Effect (see methods and Table S3 for details). (**h**) Spatial organization of the PSD-95 units based on the G-function compared to Monte-Carlo simulations.

## Discussion

Understanding homeostatic plasticity processes such as synaptic scaling requires knowledge of the complex underlying molecular mechanisms at individual synapses. Here, we focus on the scaffolding protein PSD-95, which plays a key role in regulating the positioning of neurotransmitter receptors at the postsynapse (*36*). By leveraging different labeling strategies combined with optical nanoscopy techniques, we found that prolonged neuronal activity modulation impacts the incorporation of PSD-95 at individual synaptic sites, albeit minimal nanoscale changes in the arrangement of molecules were detected.

To aid in the interpretation of the data, we would first like to discuss some relevant technical aspects of this study. To investigate whether and where newly translated PSD-95 protein is incorporated at the synapse we combined pulse-chase labeling with the ORANGE gene editing system to tag endogenous PSD-95 with HaloTag (*23*) and avoid overexpression artefacts. A similar system was recently applied to study the turnover of PSD-95 *in vivo* (*21, 22*). We were able to detect newly translated PSD-95 together with a pre-existing pool of PSD-95 over the course of 3 days. Within this timeframe we expected measurable fractions of old and new protein, since the lifetime of PSD-95 has been reported to be around 3.5 days for both native and HaloTag tagged protein (*9, 21, 37*). Yet, it is possible that not all PSD-95 molecules were visualized for two reasons. The first is incomplete labeling with the fluorescent substrates. Incorporation of the HaloTag substrate is generally a highly efficient process (*38*). However, different fluorophores exhibit different kinetics (*39*). To minimize these issues, we incubated the cells at comparably high substrate concentrations for long times (1 µM for 2 h, = 5 times higher and 8 times longer than in (*39*)) and could demonstrate that virtually all matured and accessible molecules were labeled. The second reason is the efficiency of the genome-editing process. Our cultures might contain both homozygous and heterozygous PSD-95-HaloTag knock-in cells. Therefore, we cannot exclude the presence of native PSD-95 at the analyzed synapses. Nevertheless, both aspects affect samples and conditions equally, and are therefore expected to have minimal impact on the relative quantifications performed.

Another relevant technical aspect of the study is the use of 3D MINFLUX nanoscopy to study the spatial arrangement of nanobody-labeled PSD-95. Compared to conventional antibodies, nanobodies have the advantage of minimizing the linkage error (*40, 41*), abolishing the risk of artificial protein clustering, and ensuring a higher degree of labeling of the structures of interest due to the reduced steric hindrance (*28, 32*). However, whether all PSD-95 molecules are successfully decorated by a nanobody cannot be reliably tested, since synaptic sites lack a predictable geometry and stoichiometry. Still, estimates of the labeling efficiency can be made based on other microscopic and biochemical data. When comparing our median number of 102 TIDs (*i.e.* individually detected molecules) and 49 units (*i.e.* group of TIDs that may correspond the same molecule measured repeatedly) per PSD-95 cluster with proteomic data (300-500 PSD-95 molecules per synapse, (*42, 43*)), we can assume a conservative detection rate of around 10-30%. However, an important aspect to be consider, is the stringent analysis applied (DBSCAN with a search radius of 75 nm), which on the one hand excludes a large number of molecules present at the synapse which are not incorporated into the postsynaptic density, and on the other hand can identify multiple clusters in a single dendritic spine (**Fig. S4**). When performing a less stringent analysis (**Table S1**, search radius of 200 nm), 268 TIDs and 117 units per cluster were detected, which would correspond to a nearly 90% detection efficiency in the most optimistic scenario. Ultimately, an exact quantification of the detection efficiency and protein numbers requires the use of different reporters, such as photoactivatable PaX dyes (*44*), or extensive sequential imaging rounds (*45*). However, these tools have not yet been tested for labeling crowded compartments as synaptic sites in MINFLUX experiments. A last technical consideration on the MINFLUX measurements, is that despite an active stabilization of the sample, residual drift can appear during long measuring times. Our data has been corrected with redundant cross-correlation (RCR) (*46*), the state-of-the-art approach for STORM data. However, due to the lack of a ground truth, the corrections may be affected by RCR constraints. The implementation of further hardware (*e.g.* monitoring of beam position) and postprocessing approaches would enable measurements with higher precision.

Having dissected the technical limitations, we will now discuss the biological outcome of the study, starting with the observations under basal conditions. The pulse-chase experiments indicate a higher integration of new PSD-95 protein at larger synapses compared to small synapses, however the ratio between old and new PSD-95 remained constant throughout the entire synaptic population. This data, which is consistent with observation at the presynaptic level (*47*), suggests that the process of incorporating new proteins is regulated at an individual synaptic level rather than at the cellular level. Potential mechanisms through which such synaptic specificity is achieved comprise the targeted transport to individual synapses based on synaptic tagging or an increased availability of protein production machinery in the proximity of larger synapses (*25, 48–51*). Further work is required to clarify whether the incomplete mingling of old and new PSD-95 protein pools has a functional meaning, and whether it applies to both proximal and distal dendrites (*21*). While STED provided temporal information on protein translation at synaptic sites at the resolution of molecular ensembles, 3D MINFLUX allowed to study the organization of PSD-95 at the near-molecular level. With this technique, we observed a non-random, dispersed (*i.e.* structured with a preferred distance) organization of the PSD-95 in the majority of the synaptic clusters. The data is in agreement with the observation of a retiolum-like structure by ONE expansion microscopy (*20*) and with previous observations in neurons derived from PSD-95-HaloTag knock-in animals, in which 3D MINFLUX showed a regular clustering of localizations below the <30 nm range (*19*). It is also a step forward from and complementary to the previously reported concept of nanodomains (*17, 18, 52*). In fact, the analyzed PSD-95 clusters, identified by using a search radius of 75 nm, can be considered as individual nanodomains, whose reported lower boundary is ∼80 nm. Therefore, the presented data can be considered as a zoom in the nanodomain structure. In addition to its dispersed nature, the PSD-95 organization within clusters is characterized by its robustness and independence from cluster size. While a strong correlation was observed between the volume of the cluster and the number of PSD-95 units, the distance between the units showed only a negligible dependence on this parameter. This data, together with the evidence of the partial segregation of the old and new protein pools, confirms the hypothesis of a modular organization of the PSD-95 scaffold. By adding and removing these modules, changes to the postsynaptic structure can be accomplished (*12, 53*).

Moving on to discuss the effect of long-term activity modulation, the minimal changes observed in the PSD-95 organization upon long-lasting inhibition or enhancement of neuronal activity further support the idea of a structure that is robust to changes. However, the data does not rule out the possibility of fast, immediate changes (*14*), nor of changes at the molecular level. Hints for such molecular changes are the differences in nearest neighbor median distance at the TID level and the higher number of clusters with a dispersed organization in TTX-treated samples. Further work and improvements of the currently available labeling strategies are required to achieve the actual molecular resolution and distinguish individual PSD-95 monomers, dimers, or clusters. Nevertheless, this data indicates that the organization of synaptic sites during scaling reaches a stable status comparable to the basal conditions, albeit the size and number of nanodomains in the synapse might change (*17*). Notably, while the organization of PSD-95 clusters at the nanoscale is rather robust, the modulation of the protein turnover is affected by activity changes. In particular, prolonged neuronal silencing with TTX increased the ratio of newly synthesized PSD-95, while activity enhancement with GZ had the opposite effect. These treatments linearly affected all synapses depending on their size. The elevated deposition of new PSD-95 could be a contributing factor to synaptic scaling, as already observed upon enduring TTX treatment in other studies (*26, 54, 55*). On the other hand, the reduced ratio of new over old PSD-95 induced by activity enhancement could be caused by a stalling of the protein synthesis. Regardless of the activity status, how newly synthesized PSD-95 is actually integrated and incorporated into the robust scaffold is unclear. This question could be answered by – still challenging – two-color 3D MINFLUX measurements of pulse-chased cultures. These experiments, if performed under live conditions, would also rule out fixation artifacts.

In conclusion we have combined different imaging methods to quantify the behavior of PSD-95 at the postsynapse as a function of neuronal activity at nanoscale and near-molecular resolution. STED analysis of hundreds of synapses showed that neuronal activity affects the rate of protein synthesis at the postsynapse. MINFLUX nanoscopy allowed to analyze PSD-95 organization at unprecedented resolution, revealing a non-random, dispersed, structure which is largely robust and independent of synaptic size and long-term activity changes. Altogether we show that postsynaptic densities have a relatively fixed but still dynamic structure, capable to incorporate newly synthesized protein according to the specific synaptic needs.

## Materials and Methods

### Preparation of neuronal cultures

All procedures were performed in accordance with the Animal Welfare Act of the Federal Republic of Germany (Tierschutzgesetz der Bundesrepublik Deutschland, TierSchG) and the Animal Welfare Laboratory Animal Regulations (Tierschutzversuchstierverordnung) and no specific authorization or notification was required. The procedure for sacrificing P0–P2 rats performed was supervised by animal welfare officers of the Max Planck Institute for Medical Research (MPImF) and conducted and documented according to the guidelines of the TierSchG (permit number assigned by the MPImF: MPI/T-35/18).

Primary hippocampal cultures were prepared from dissociated tissue of P0-P2 postnatal wild-type Wistar rats of either sex (Janvier-Labs) as described previously in D’Este et al. 2015; Gürth et al. 2020. Briefly, dissected hippocampal tissue was digested with 0.25% trypsin for 20 minutes at 37°C, mechanically dissociated by pipetting, and maintained in Neurobasal supplemented with 2% B27, 1% GlutaMAX and 1% penicillin/streptomycin (all from Gibco, Thermo Fisher Scientific). Cells were seeded at a concentration of 110,000/well in 12-well plates on Ø 18mm glass coverslips coated with 0.1 mg/ml poly-ornithine (Sigma) and 1 µg/ml laminin (Corning). Medium was changed to fresh supplemented Neurobasal 1-2 hours after seeding and cultures were maintained in an incubator (37°C, 5%CO2) until DIV21-25 in presence of residual glial cells.

### Plasmid generation and viral vector production

Plasmids pAAV-hSyn-PSD-95-HaloTag for overexpression controls was constructed as described in (Masch et al. 2018), pAAV-EFS-SpCas9 (Addgene #104588) was a gift from Ryohei Yasuda (Nishiyama et al. 2017), and pAAV ORANGE Dlg4-HaloTag KI (Addgene #139656) was a gift from Harold MacGillavry (Willems et al. 2020). The preparation of adeno-associated viral vectors was performed as previously described in Masch et al. 2018 and purified as described in Zolotukhin et al. 2002.

### Pulse-chase labeling of PSD-95-HaloTag

Cultured neurons were infected at DIV7 with AAV-ORANGE-Dlg4-Halo (PSD-95) and AAV1/2-EFS-SpCas9 or rAAV1/2-hSyn-PSD-95-Halo. Mature cultures (DIV15-20) were labeled with 1 µM 580CP-Halo (pulse) in preconditioned Neurobasal for 2 hours at 37°C to achieve complete labeling of tagged structures. Afterwards cultures were washed in warm ACSF (126 mM NaCl, 2.5 mM KCl, 2.5 mM CaCl_2_, 1.3 mM MgCl_2_, 30 mM Glucose, 27 mM HEPES) and brought back in culture in their own preconditioned Neurobasal.

After further 48 hours in culture the labeled samples were relabeled with the SiR-Halo (chase) dye at 0.5 µM for 24h at 37°C in preconditioned Neurobasal. For testing the influence of neuronal activity, concomitantly to the chase application, cultures were stimulated with 1.5 µM Tetrodotoxin (TTX, Tocris, cat. 1078) or 10 µM Gabazine (GZ, SR 95531 hydrobromide, Tocris, cat. 1262).

Prior to immunolabeling cultures were fixed in 4% PFA in PBS, pH 7.4, quenched with quenching buffer (PBS, 100 mM glycine, 100 mM ammonium chloride), permeabilized for 5 min in 0.1% Triton X-100 and blocked with 1% BSA for 1 hour. Samples were then incubated for 1-2 hours at room temperature each with primary antibodies, nanobodies, secondary antibodies and dyes, each followed by five washes in PBS. Antibodies, nanobodies, and dyes used were anti-Halo-Tag (Promega, cat. G928A, 1:200 dilution), FluoTag-X2-anti-PSD95-Sulfo-Cy5 (*28*)(NanoTag Biotechnologies GmbH, cat. N3702-SC5-L, 1:200 dilution), FluoTag-X2-anti-PSD95-580 (NanoTag Biotechnologies GmbH, cat. N3702-AbStar580-L, 1:200 dilution), Alexa Fluor 405 anti-rabbit (Thermo Fisher, cat. A31556, 1:100 dilution), Phalloidin Alexa Fluor 405 (Thermo Fisher, cat. A30104; 1:200 dilution), and Phalloidin StarGREEN (Abberior GmbH, cat. STGREEN-0100-20ug, 1:100 dilution). Samples for confocal and STED imaging were afterward embedded in Mowiol® supplemented with DABCO and cured for at least 1 hour at RT before imaging.

### PSD-95 nanobody purification and conjugation to DNA strand

For site specific coupling of nanobodies, the procedure from Schlichthaerle et al. was modified (*56*). Briefly, unconjugated anti-PSD-95 nanobodies (NanoTag Biotechnologies GmbH) carrying an ectopic cysteine on the C-terminus allow chemical coupling with a maleimide functional group on the DBCO-maleimide crosslinker (Sigma-Aldrich, cat. 760668) and finally to the P2 (5’-TATGTAGATC-3’) docking DNA-strands (Biomers GmbH, Ulm, Germany). Therefore, the nanobody ectopic cysteine residues were reduced with 5 mM tris(2-carboxyethyl)phosphine (TCEP) for 30 minutes at 4°C. Excess TCEP was removed via 10 kDa MWCO centrifugal filters (Merck, cat. UFC501096). The DBCO-Maleimide crosslinker was added at 20x molar excess under gentle shaking for 4 h at 4°C. Excess of the crosslinker was removed using 10 kDa MWCO centrifugal filers. Then, 5x molar excess of single-stranded DNA (ssDNA), was added for 1 h at RT. The ssDNA sequence (P2), was functionalized on the 5’-end with an azide (N3) group to perform cupper-free click chemistry to the DBCO-functionalized nanobody. The ssDNA conjugated nanobody was purified using size exclusion chromatography on a GE Äkta pure 25 system equipped with a Superdex 75 increase 10/300 GL column (Cytiva, cat. 29148721).

### Sample preparation for MINFLUX

For MINFLUX experiments, hippocampal primary neurons (HPN, DIV 18-20) were treated, fixed, quenched and permeabilized as described above. Samples were then blocked in BSA 1% for 30 min. DNA-PAINT samples were incubated for 1 hour at room temperature with an anti-bassoon guinea pig antibody (Synaptic Systems cat. 141 004, 1:1000 dilution) as counterstaining and, if applicable, an anti-PSD-95 antibody (Synaptic Systems, cat. 124 011, 1:500 dilution). After washing in PBS, samples were incubated for 1 hour in the antibody incubation buffer (Massive Photonics, Buffer kit) with anti-guinea pig AlexaFluor 405 (Abcam, cat ab175678, 1:1000 dilution), phalloidin AlexFluor 488 (Thermo Fisher, cat. 12379, 1:1000 dilution), and either the anti-PSD95 nanobody conjugated to the DNA strand (final concentration ∼40 µM) or anti-mouse secondary nanobody decorated with P1 DNA strand (Massive Photonics, Massive-sdAb 1-PLEX, 1:250 dilution). Nanobodies were applied right before imaging. After washing in washing buffer (Massive Photonics, Buffer kit), gold nanorods fiducials were added to the sample (Nanopartz, cat. A12-40-980-CTAB-DIH-1-25, 1:1 in PBS for 5 min), washed and mounted on concave coverslides in high salt buffer (PBS supplemented with 500mM NaCl) and 1 nM P2 DNA-PAINT imager (TATGTAGATC) conjugated at the 3’ end to Atto655 (Biomers GmbH, Ulm, Germany), or in imaging buffer (Massive Photonics, Buffer kit) and 1 nM P1 DNA-PAINT imager conjugated to Atto655 (Massive Photonics, sdAb 1-PLEX).

For samples with sulfoCy5, coverslips were incubated for 1 hour at room temperature with FluoTag-X2-anti-PSD95-Sulfo-Cy5 (NanoTag Biotechnologies GmbH, cat. N3702-SC5-L, 1:200 dilution), and phalloidin StarGREEN (Abberior GmbH, cat. STGREEN-0100-20 µg, 1:1000 dilution). After washing, samples were mounted with gold nanorods (Nanopartz, cat. A12-40-980-CTAB-DIH-1-25) in blinking buffer (cysteamine concentration 10 mM) as described in (*57*). All MINFLUX samples were imaged within 4 days from the fixation.

### Hexanediol treatment

1,6-Hexanediol (Aldrich, cat. 240117-50G) was used at 10% w/v in preconditioned Neurobasal supplemented with 2% B27, 1% GlutaMAX and 1% penicillin/streptomycin (Hex-medium, all reagents from Gibco, Thermo Fisher Scientific, Waltham, MA, USA). The Hex-medium was kept in a semi-open falcon tube in the neuronal culture incubator for at least 30 minutes before the treatment. Cell culture media was removed, under sterile conditions, from the neuronal culture and replaced with an equal volume of 10% Hex-medium. HPNs (DIV 18-20) were incubated for exactly 2 minutes (37 ◦C, 5% CO2, 95% rH). Afterward, Hex-medium was removed and HPNs were directly fixed in 4% PFA in PBS for 20 minutes, at pH 7.4, quenched with quenching buffer (PBS, 100 mM ammonium chloride, 100 mM glycine), permeabilized for 5 min in 0.1% Triton X-100 in PBS, and blocked with 1% BSA in PBS for 30 minutes. Samples were incubated for 1 h at room temperature with primary antibodies/nanobodies and secondary antibodies in PBS, while for DNA-PAINT single-domain antibody, an incubation buffer (Massive Photonics) was used, each followed by 3 washes in PBS or washing buffer (Massive Photonics) for 5 minutes each. Homer1 rabbit (Synaptic Systems, cat. 160 003, 1:500 dilution), Bassoon guinea pig (Synaptic Systems cat. 141004, 1:1000 dilution), FluoTag®-X2 anti-PSD95 Star635P (NanoTag Biotechnologies, cat. N3702-Ab635P-L, 1:250), FluoTag® anti-PSD95 – with P2 docking strand, phalloidin STARGREEN (Abberior, cat. STGREEN-0100–20 µg, 1:500 dilution), phalloidin Alexa488 (ThermoFisher, cat. A12379, 1:1000 dilution), STAR 580 anti-rabbit (Abberior, cat. ST580-1002–500 µg, 1:500 dilution), Alexa Fluor 405 anti-guinea pig (Abcam, Cambridge, United Kingdom, ab175678, 1:1000 dilution) were used as primary antibodies/nanobodies and secondary antibodies.

Control samples for confocal imaging were embedded in Mowiol® (Sigma-Aldrich/Merck, cat. 81318) supplemented with DABCO (Sigma-Aldrich/Merck, cat. 290734) and cured for at least 3 h at room temperature before imaging. DNA-PAINT samples for MINFLUX were further processed as described above.

### Confocal, STED and MINFLUX imaging

Samples were imaged on an Abberior Expert Line Microscope (Abberior Instruments GmbH) on a motorized inverted microscope IX83 (Olympus) and equipped with pulsed STED lines at 775 nm and 595 nm, excitation lasers at 355 nm, 405 nm, 485 nm, 580 nm, and 640 nm, and spectral detection. Spectral detection was performed with avalanche photodiodes (APD) and detection windows were set to 650-725 nm, 600-630 nm, 505-540 nm, and 420-475 nm to detect Atto647N/Star635P, Alexa Fluor 594/Star580, Alexa Fluor 488/StarGREEN and Alexa Fluor 405, respectively. Images were acquired either with a 20x/0.4 NA oil immersion lens with pixel size at 100 nm or with a 100x/1.4 NA oil immersion lens, 30 nm pixel size, and pinhole to 100 µm (1 AU). Laser powers and dwell times were kept consistent in all experiments.

MINFLUX Imaging was performed on an Abberior 3D MINFLUX (Abberior Instruments GmbH) built on a motorized inverted microscope IX83 (Olympus) and equipped with 640 nm, 561 nm, 485 nm, and 405 nm laser lines, and active sample stabilization with an IR laser (975 nm). Detection was performed with 2 APDs in the spectral windows 650-685 nm and 685-720 nm. Images were acquired using the default 3D imaging sequence, with an L in the last iteration step of 40 nm, and photon limit of 100 and 50 photons for the lateral and axial localization, respectively (**Table 1**). Power at the periscope in the first iteration step was ∼330 µW for DNA-PAINT experiments (pinhole 0.8 A.U. corresponding to 60 µm) and ∼80 µW for sulfoCy5 experiments (pinhole 0.93 A.U. corresponding to 70 µm). Postsynaptic sites were identified based on the phalloidin staining and either the PSD-95 signal in the case of sulfoCy5, or bassoon for the DNA-PAINT samples. Multiple regions of interest within a 15 x 15 µm field of view were selected for imaging. Image acquisition of sulfoCy5 samples was interrupted when the number of localizations was not increasing upon higher UV activation. Measurements of DNA-PAINT image acquisition lasted from a minimum of 3 hours up to overnight.

**Table 1.** Sequence utilized for 3D MINFLUX imaging. Parameters utilized for each iteration step of the MINFLUX imaging procedure are provided.

| Iteration | Modality | L (nm) | Photon limit | Background limit (Hz) | Pattern dwell time (ms) | Pattern repeat | Centre Frequency Ratio limit | Laser power factor |
| --- | --- | --- | --- | --- | --- | --- | --- | --- |
| 0 | Confocal | - | 160 | 15000 | 1 | 1 | - | 1 |
| 1 | z-line | - | 400 | 15000 | 1 | 1 | - | 1 |
| 2 | Square | 288 | 100 | 10000 | 1 | 5 | 0.8 | 1 |
| 3 | z-line2 | 288 | 50 | 10000 | 1 | 5 | - | 1 |
| 4 | Square | 151 | 67 | 10000 | 1 | 5 | - | 2 |
| 5 | z-line2 | 151 | 33 | 10000 | 1 | 5 | - | 2 |
| 6 | Square | 76 | 67 | 10000 | 1 | 5 | - | 4 |
| 7 | z-line2 | 76 | 33 | 10000 | 1 | 5 | 0.8 | 4 |
| 8 | Square | 40 | 100 | 10000 | 1 | 5 | - | 6 |
| 9 | z-line2 | 40 | 50 | 10000 | 1 | 5 | - | 6 |

### Processing and analysis of STED data

Images shown in figures were visualized and processed with Imspector (Abberior Instruments GmbH) and FIJI (*58*). STED images are shown as raw data. Brightness was adjusted uniformly throughout the images. MINFLUX data was analyzed in Matlab (version 2021b, MathWorks) and Python (Python Software Foundation).

To quantify the relative amount of PSD-95-Halo in STED images, PSD-ROIs were determined based on the confocal 488 nm channel, in which the anti-HaloTag antibody was imaged. Therefore, “moments” threshold was applied and background signal (below 0.2 µm²) was eliminated with the analyze particles function. The ROIs were then used to determine signal area and intensity in the 640 nm and 580 nm STED channels by applying “moments” or “isodata” thresholding, respectively and then measuring area and intensity within this thresholded area for each individual PSD site separately. The protein amount was calculated by multiplying the signal area by the mean signal intensity and normalized with respect to the control and is expressed in arbitrary units (A.U.). For images displaying imaging artefacts that influenced the overall maximum intensity (reflections or dye accumulations resulting in bright non-specific puncta) a maximum intensity was individually set and exceeding extreme values were cropped to the default maximum using Matlab. The mean amounts of old and new proteins in treated samples were normalized for each respective experimental round to the mean of DMSO treated samples (**Fig. 2b**). Within each treatment condition, the amounts for each experimental round were binned into 5 equally spaced intervals and the respective mean values for each bin were fitted with a linear regression (**Fig. 2d-f**), similarly of what performed in (*24, 25*).

For colocalization analysis, PSD-95 signal was segmented from the background using a binary mask. The mask was generated by saturating 99% of the original image, followed by an 8 pixels median filtering and an automatic Otsu’s global threshold. ROIs smaller than 200 pixels or greater than 25000 pixels were excluded from the binary mask, in order to exclude background noise or dye accumulation. The binary mask was then applied to each respective image, and the Pearson correlation coefficient and the Manders coefficients were computed. The former coefficient considers both cooccurrence and intensity of the signal (*i.e.* colocalization), and the latter considers cooccurrence only. The analysis was made using a custom MATLAB code.

### Processing and analysis of confocal data

The relative amount of homer in PSDs with respect to dendrites ROIs was quantified by computing the mean intensity of the homer channel after applying binary masks to PSD and dendrites regions. The PSD masks were generated after applying a threshold at the 1% highest intensities followed by image disk dilation with 2 pixels. The dendrites masks were generated after applying a multi threshold Otsu segmentation with 3 levels. The lowest level was assigned to background and the complement to dendrites. Then, the intersection with PSD ROIs were removed. Afterwards, the mean intensity of homer channels masked with PSD ROIs and dendrites ROIs was computed.

### MINFLUX data analysis

Prior to analysis, the data was drift corrected using the redundant cross-correlation algorithm implemented in Matlab as in (*46*). Localizations from the same emission trace, i.e. with same trace identification (TID), farther than three standard deviations with respect to the mean trace position were considered outliers and excluded from the trace. Only the traces containing at least 4 localizations were considered. The experimental localization precision was estimated by computing the median value of the trace standard deviation (*59*).

The point pattern spatial analysis was done considering the TID centroid position as a point in a 3D Euclidian space. A density-based spatial clustering of applications with noise (DBSCAN) algorithm was applied to define the 3D PSD Clusters. The MATLAB built-in function *dbscan* was used with a minimum number of neighbors *(minPoints*) equal to 40 and a search radius (*eps*) of 75 nm, unless otherwise stated. For the PSD units clustering, another DBSCAN was performed with *minPoints* = 1 and *eps* = 8.5 nm. For each PSD cluster, the 3D alpha shape was retrieved using the MATLAB *alphashape* function with the default critical alpha radius. The PSD volume, superficial area and x-y projected area were derived from each cluster alpha shape. The compactness was defined as the cubic sphericity, *i.e. π x 36 x V²/S²* where *V* is the volume and *S* the superficial area. To test for spatial randomness (SR) of PSD Units distribution within the PSD clusters, the Diggle-Cressie-Loosmore-Ford (DCLF) test (*33*) was applied using Monte Carlo simulations with the G-function as spatial descriptor. For the Monte Carlo Simulations, each synthetic PSD cluster consisted of randomly distributed points, conditioned by the number of PSD Units, the minimum distance of 8.5 nm and the volume of the observed PSD clusters. For each observed cluster, 1000 Haloetic clusters under SR were generated. The observed G function (G_OBS_) was then, compared with the expected G function under SR (G_SR_) and its 95% confidence envelope. Afterwards the DCLF test was applied considering the full range of nearest neighbor distances. All the simulations and tests were done with a custom code written in MATLAB. A visual representation of the data processing is provided in **Fig. S4**.

### Statistical analysis

To quantify the colocalization of old and new protein compared to multicolor fluorescent beads and control image, the Pearson non-zero coefficient was applied based on pixel intensity. Mean and SEM values of 0.8030 ± 0.05 and 0.4832 ± 0.0238 for control (N = 4) and PSD old-new (N=15). Correlation of old versus new protein amounts at individual synaptic sites (n=2148, N=5) was quantified using Spearman’s correlation coefficient (ρ= 0.823) (**Fig 1 e-f**). Differences of relative overall protein turnover was tested using analysis of variance followed by Dunnet multiple comparison correction. Mean and SEM_N=4_ values were: TTX_old_ = 1.23 ± 0.10, p = 0.22, TTX_new_ = 1.74± 0.15, p = 0.03 and GZ_old_ = 1.15 ± 0.11, p = 0.47, GZ_new_ = 0.80± 0.03, p = 0.01 (**Fig 2** **b)**. To interpret the activity dependence of PSD-95 organization considering the grouping effects of experimental replicates and PSD-95 clusters belonging to the same image, the data was modelled using Linear Mixed Model Effect. Maximum likelihood estimation was used as fitting method. Treatments were modelled as fixed effects and the experimental rounds and images as random effects. For each parameter of interest, the best model was chosen according to Akaike Information Criterion. Detailed information about the chosen model, the estimates, 95% confidence interval and p-values are described in Supplementary Information (**Table A1**). In **Fig. 4** **h**, Chi-squared test was used to compare the proportions of random and non-random PSD clusters treated with TTX and GZ against the expected values of cells treated with DMSO: Χ_TTX_ = 8.089, p = 0.0045 and Χ_GZ_ = 0.150, p = 0.6984. Unpaired t-test was used to compare Hex treated cells versus control. Control(_N=4)_ = 3.11 ± 0.32, Hex_(N=8)_ = 1.93 ± 0.20, t(_12_) =7.92, p = 0.00001 (**Fig. S8)**.

## Acknowledgments

We thank Prof. Dr. Stefan W. Hell for supporting the project. We thank the MPI-NAT chemistry facility for the synthesis of HaloTag substrates. We thank Eugenio F. Fornasiero and Birgit Koch for support with gene editing tools. We thank Annette Herold, Magnus-Carsten Huppertz, Birgit Koch, Jana Kress and Alena Fischer for support with neuronal cultures and virus production. We thank Lorenzo Bergamo for critical reading of the manuscript.

## Author contributions

Conceptualization: CMG, ED

Methodology: CMG, MADRBFL, JH, ARCD

Investigation: CMG, VMP, JH, ED

Visualization: CMG, MADRBFL, ARCD, ED

Resources: JH, NM, FO

Formal analysis: CMG, MADRBFL

Supervision: FO, ED

Writing—original draft: CMG, MADRBFL, ED

Writing—review & editing: all authors

