## Supplementary materials for "Neuronal activity modulates the incorporation of newly translated PSD-95 into a robust structure as revealed by STED and MINFLUX"

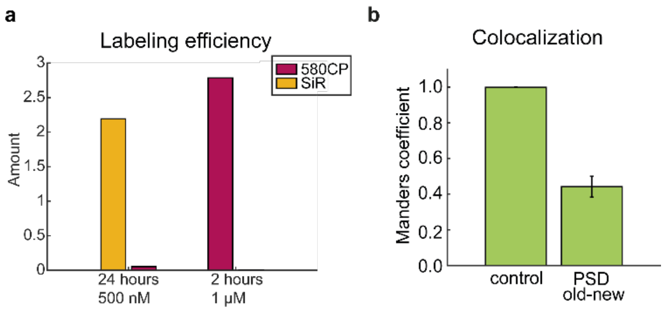


**Fig. S1. Performances of the pulse-chase labeling and dual color STED imaging.**
(**a**) Control of the complete labeling of endogenously expressed PSD-95-Halo tag is the conditions in which the pulse and chase labeling have been performed. Neurons were labeled with either SiR-Halo at 0.5 µM concentration for 24 h or with 580CP-Halo at 1 µM concentration for 2 h. This step was immediately followed by labeling with the respective other Halo substrate for 1h at 1 µM concentration. Values represent the mean relative protein amount (expressed in A.U.) calculated as the product of pixel area and mean intensity of each individual synaptic site (n = 288-644 synaptic sites from 3 images, N = 1 independent experiment). (**b**) Manders co-occurrence coefficient of old and new protein in dual color STED images compared to control (same image at different resolutions). Note that the Manders coefficient measures only the overlap of two signals, without considering their brightness. Mean and SEM values of 0.9986 ± 0.0014 and 0.4419 ± 0.0584 for control (n = 4) and PSD old new ($n_{images}$ = 15, $N_{independent\ cultures}$ = 5), respectively. Same dataset as Fig. 1e.

**a**

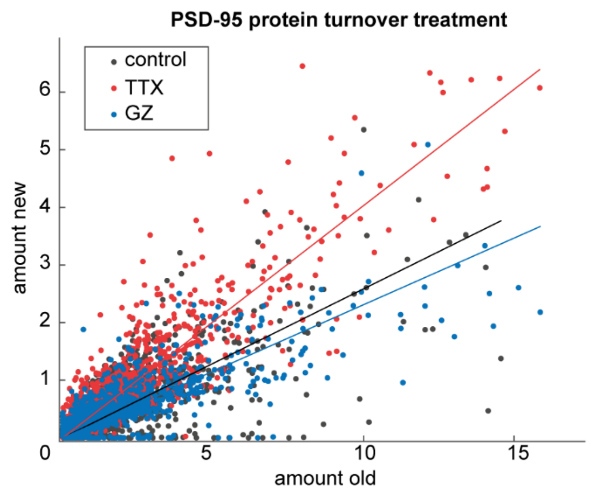


**b**

| Treatment | b | $R^2$ | ρ |
|---|---|---|---|
| **DMSO** | 0.259 | 0.478 | 0.781 |
| **TTX** | 0.403 | 0.782 | 0.882 |
| **GZ** | 0.231 | 0.611 | 0.777 |

**Fig. S2. New PSD-95 translation upon treatment – raw data.**
The data represents the not-binned data shown in Fig. 2d-f. (**a**) Correlation of the amount of old PSD-95-Halo protein and new PSD-95-Halo protein for DMSO (n = 1858), TTX (n = 1183) and GZ (n = 790) treated cultures from 4 independent experiments. The dependence of old and new protein amounts is described by the Spearman's correlation coefficient (ρ) and linear fits. (**b**) Parameters of correlation for each treatment. Data was fit with a first order polynomial curve with slope *b*. The goodness of the fit ($R^2$) describes the linear dependence of amount of old vs new protein, while the correlation was assessed by Spearman's correlation coefficient (ρ).

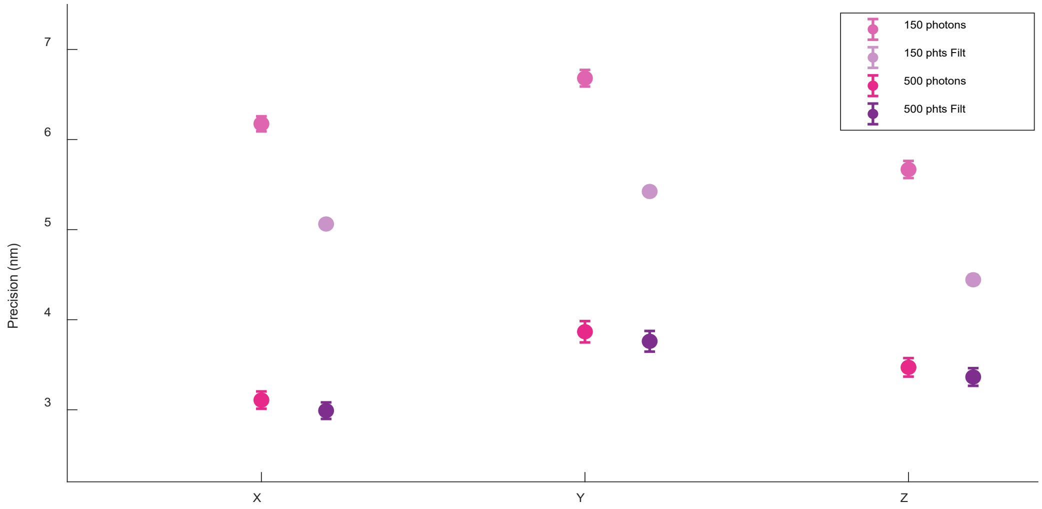


| N Photons | Condition | $\bar{X} \pm \sigma$ (nm) | $\bar{Y} \pm \sigma$ (nm) | $\bar{Z} \pm \sigma$ (nm) |
|---|---|---|---|---|
| 150 | RAW | 6.17± 0.08 | 6.68 ± 0.09 | 5.67 ± 0.10 |
| 150 | Filtered | 5.06 ± 0.05 | 5.42 ± 0.05 | 4.44 ± 0.05 |
| 500 | RAW | 3.11 ± 0.10 | 3.87 ± 0.12 | 3.47 ± 0.10 |
| 500 | Filtered | 2.99 ± 0.09 | 3.76 ± 0.11 | 3.36 ± 0.10 |

**Fig. S3. 3D MINFLUX localization precision in DNA-PAINT experiments.**
Plot with mean values for localization precision in x, y, and z, and error bars representing standard error of the mean. Results from 33 images for Raw and Filtered data, before (150 photons) and after (500 photons) photon aggregation are shown. Below, table with descriptive values.

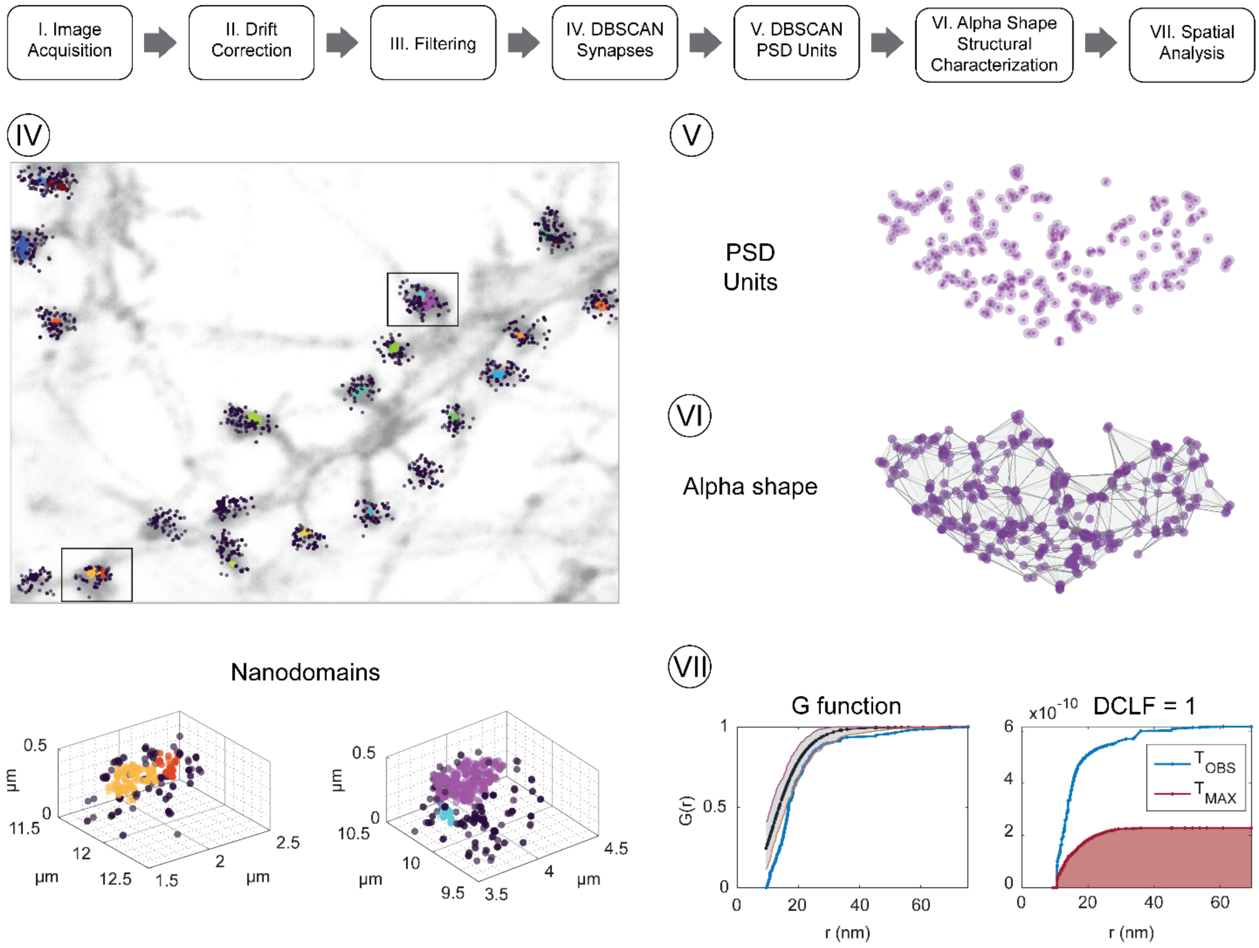


**Fig. S4. MINFLUX data processing workflow.**
MINFLUX data processing workflow with exemplary images for the steps IV-VII.
Step IV: noise removal with DBSCAN to identify the PSD-95 clusters. Confocal image of phalloidin (actin, gray) overlaid with MINFLUX TIDs rendered as gaussian with FWHM of 12 nm. After DBSCAN, PSD-95 clusters are identified. Colored TID are assigned to a PSD-95 cluster, while darker circles correspond to excluded TIDs. Each PSD-95 cluster is rendered with a different color. Note that within the same spine, multiple PSD-95 clusters can be identified (lower panels show the close-up of the spines marked with a box).
Step V: A second DBSCAN (eps = 8.5 nm) is performed on the TIDs assigned to a PSD-95 cluster to group TIDs into units. The number of objects and nearest neighbor distances are calculated.
Step VI: Individual PSD clusters after noise removal are analyzed based on the alpha shape geometry. Cluster volume, density, and compactness are calculated based on the alpha shape.
Step VII: Based on the alpha shape, the observed units distribution is compared to random particle organization as determined by Monte Carlo simulations with G function (right) and DCLF test (left).

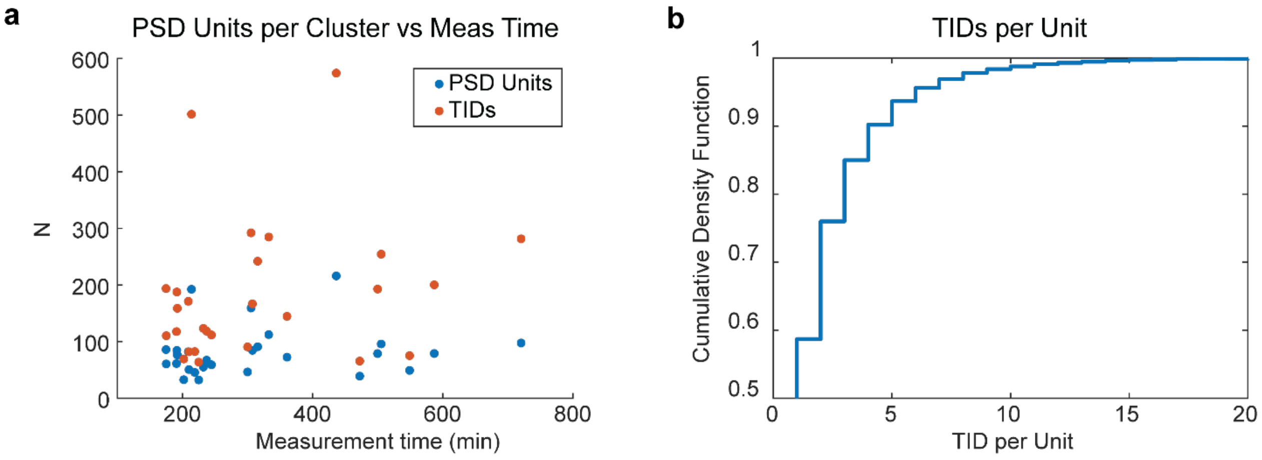


**Fig. S5. Impact of imaging time on TIDs and units in DNA-PAINT experiments and number of TIDs per unit.**

(**a**) The number (N) of units (blue) and TIDs (orange) per PSD-95 cluster does not scale linearly with the imaging time. The Pearson correlation coefficient is 0.15 with a p-value of 0.46 for the units and 0.23 with a p-value 0.24 for the TIDs, showing no significant linear correlation between the two variables. (**b**) Cumulative Density Function for the number of TIDs per PSD units in basal conditions.

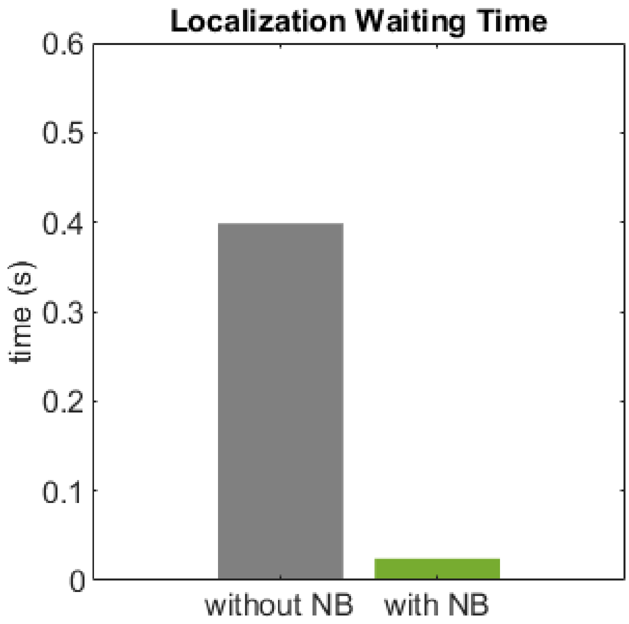


**Fig. S6. Localization waiting time for samples without and with nanobodies.**
The average time required to detect a localization in control samples without nanobody (NB) was around 0.4 seconds compared to 0.025 seconds for samples with NB.

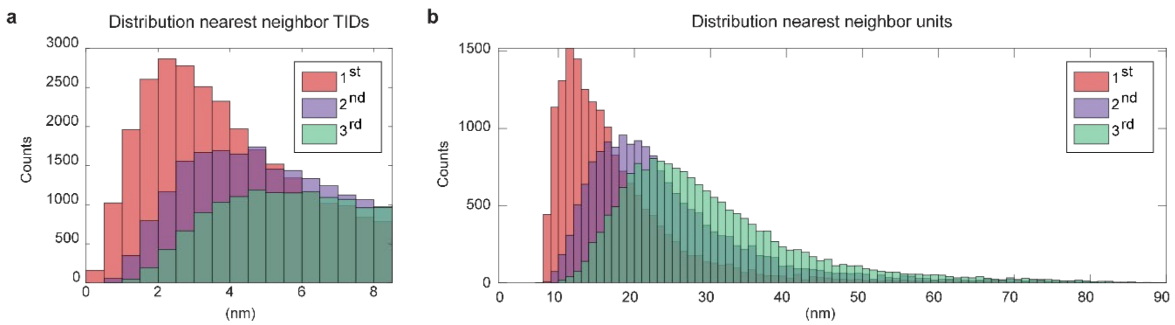


**Fig. S7. PSD-95 nearest neighbor distances characterization.**

Distribution of 1st, 2nd and 3rd nearest neighbor distances for TIDs (**a**) and PSD-95 units (**b**) in PSD-95 clusters in basal conditions. Same dataset as Fig. 3.

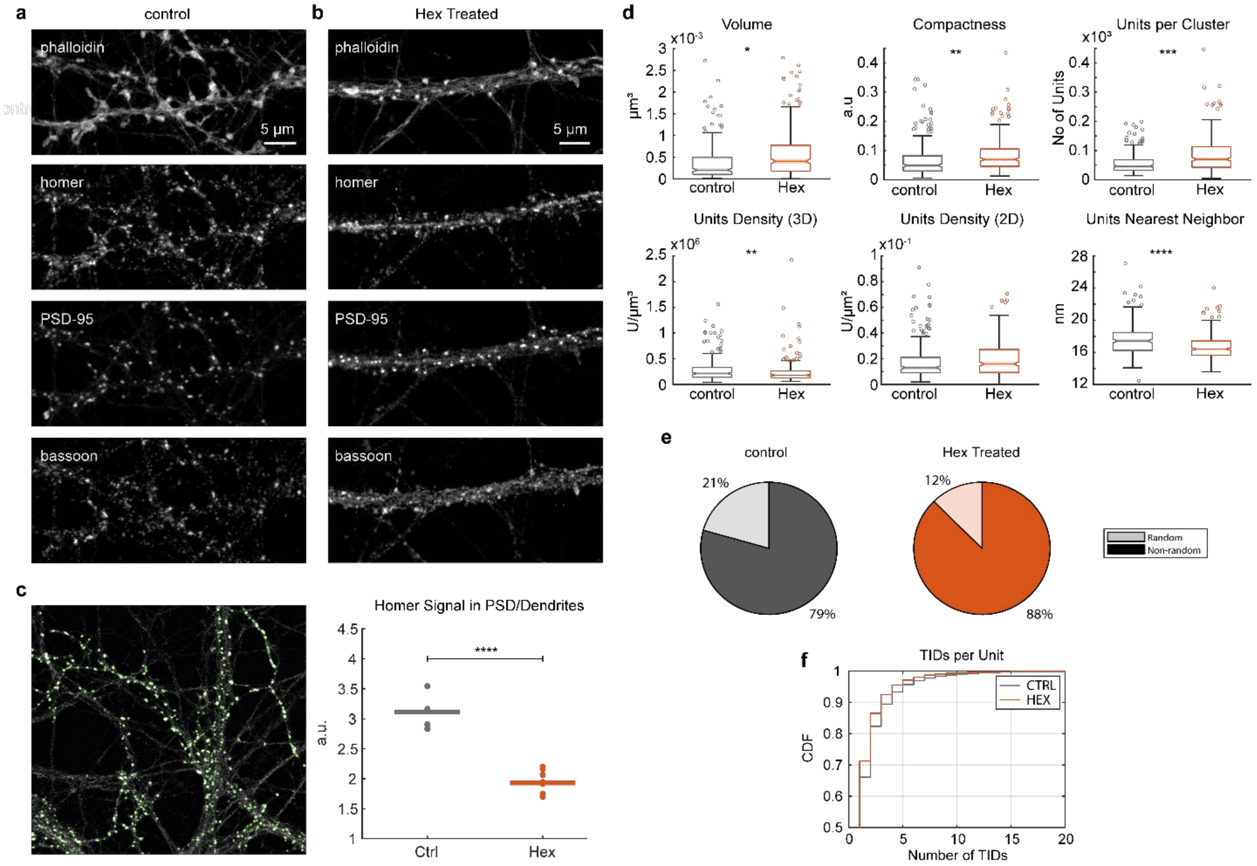


**Fig. S8. Changes in the organization of PSD-95 caused by 2 min incubation with 1,6-Hexanediol.**

Confocal images of (**a**) control or (**b**) 1,6-Hexanediol-treated neurons labeled against actin (phalloidin), homer, PSD-95 and bassoon. Scale bars: 5µm. (**c**) Example of image segmentation based on the PSD-95 signal. Green outlines indicate areas which are identified as synaptic sites. On the right, quantification of homer fluorescence intensity signal in the PSD-95 segmented areas over dendritic shafts. (**d**) Effect of 1,6-Hexanediol treatment on PSD-95 clusters (number of analyzed PSD-95 clusters: control = 243, Hex = 281 from 9-11 cells and 3 independent experimental rounds). * $p<0.05$; ** $p<0.01$; *** $p<0.001$. Statistical analysis performed using Linear Mixed Model Effect (see methods). (**e**) Percentage of synaptic sites with a random or non-random (dispersed) organization of PSD-95 units. (**f**) Cumulative distribution function of the number of TID per unit in control or Hex-treated samples.

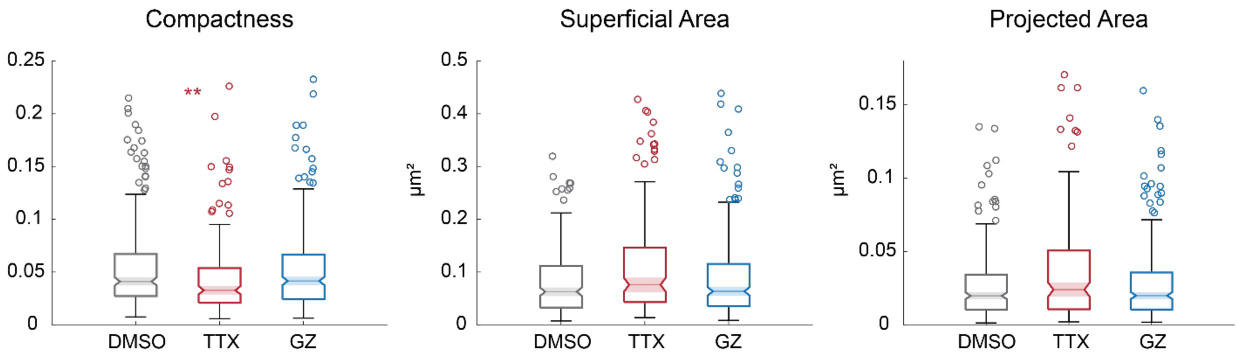


**Fig. S9. 3D MINFLUX characterization of activity dependent PSD-95 cluster morphology.** (**a**) Compactness (a.u.), (**b**) superficial area and (**c**) 2D projected area (in xy-plane) of PSD-95 clusters. Same dataset as in Fig. 4.

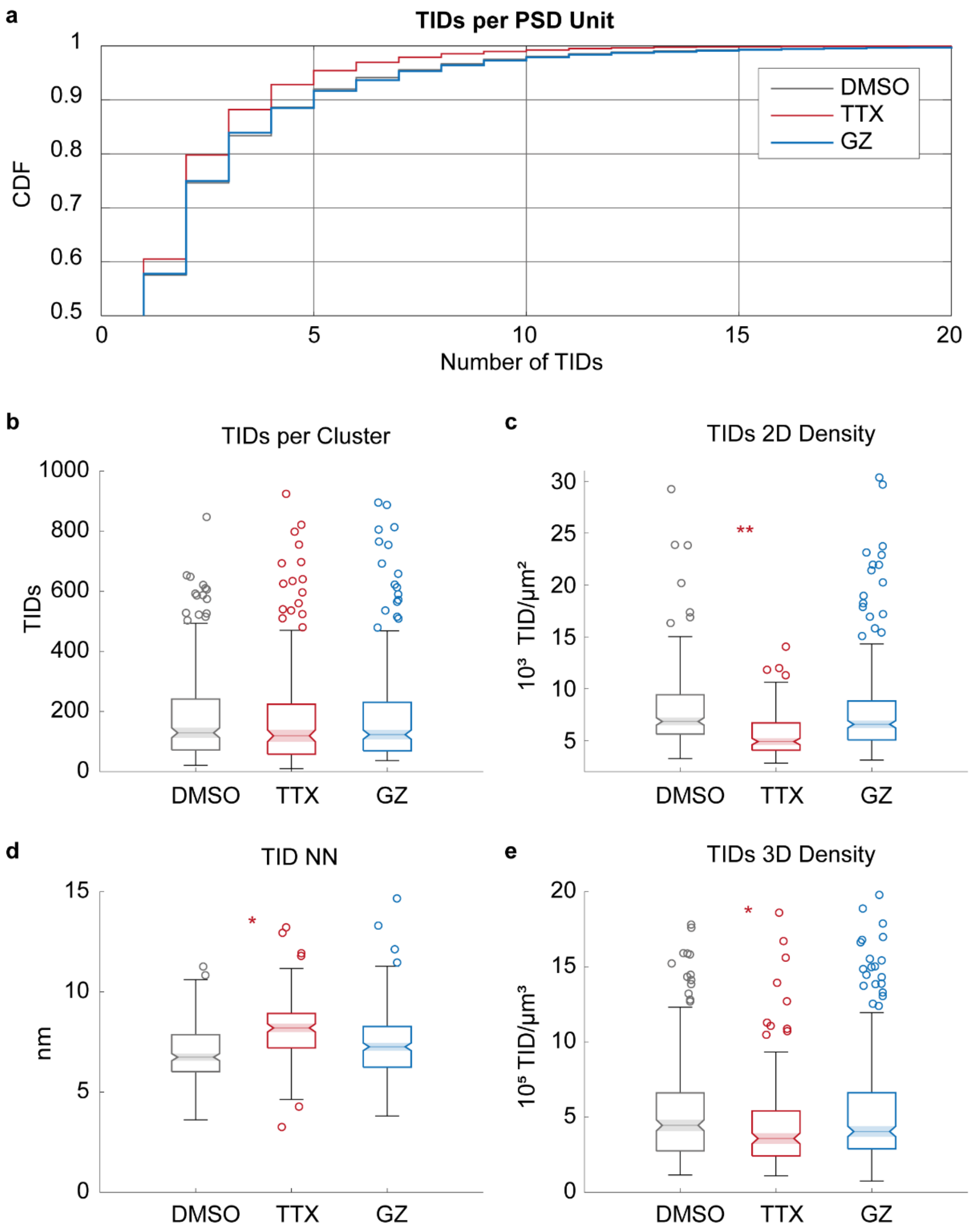


**Fig. S10. 3D MINFLUX characterization of activity dependent changes of PSD-95 organization based on trace IDs.**

(**a**) Cumulative distribution function of TIDs per PSD unit in DMSO-control, TTX, or Gabazine (GZ) treated samples. (**b**) Quantification of trace IDs per cluster, (**c**) TIDs 2D density based on the cluster projected area in xy-plane, (**d**) Nearest Neighbor distances (using Euclidian metric) between each TID, (**e**) 3D density of the TIDs depending on the treatment. Same dataset as in Fig. 4.

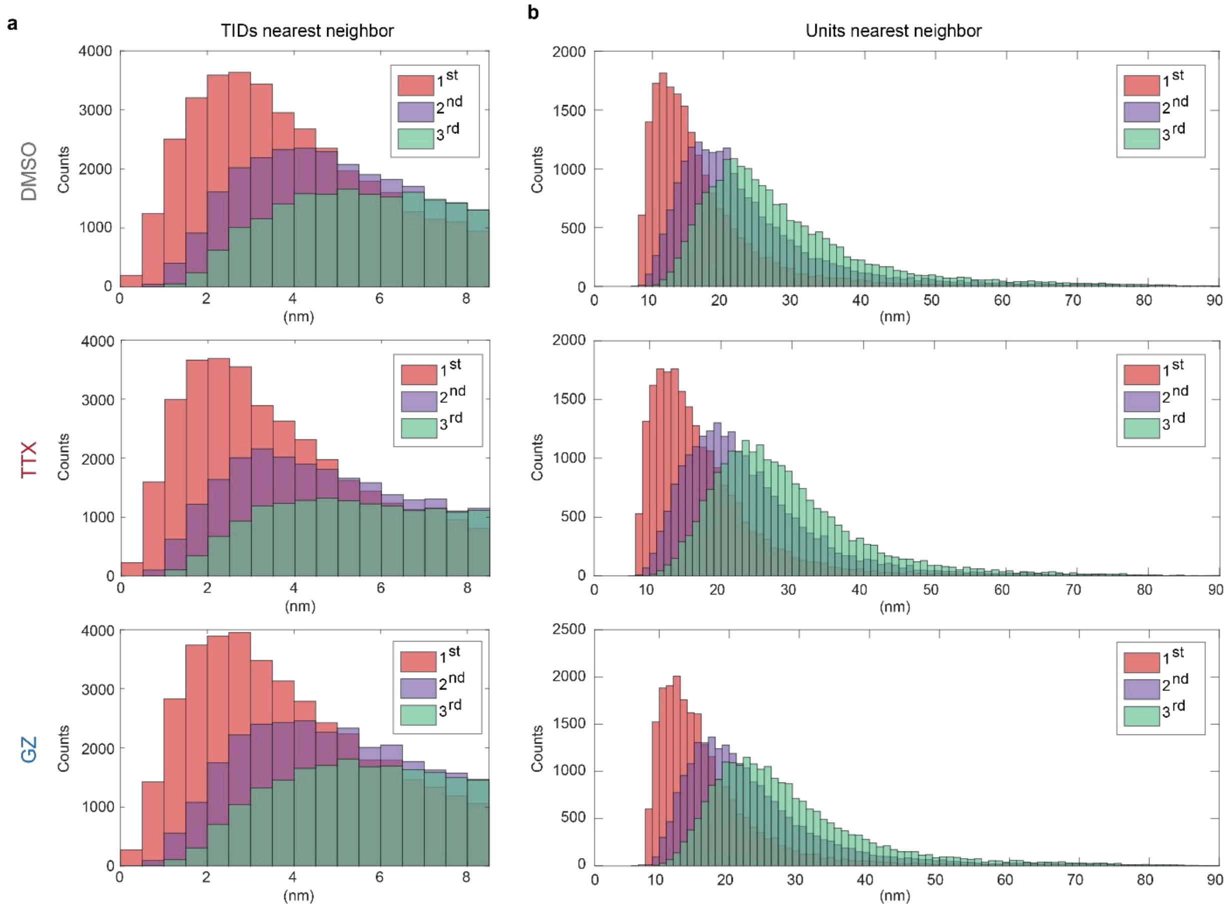


**Fig. S11. PSD-95 nearest neighbor distances of TIDs and units in treated samples.**
Distribution of the 1st, 2nd and 3rd nearest neighbors (**a**) TIDs or (**b**) units in DMSO, TTX or Gabazine (GZ) treated samples. Same dataset as in Fig. 4.

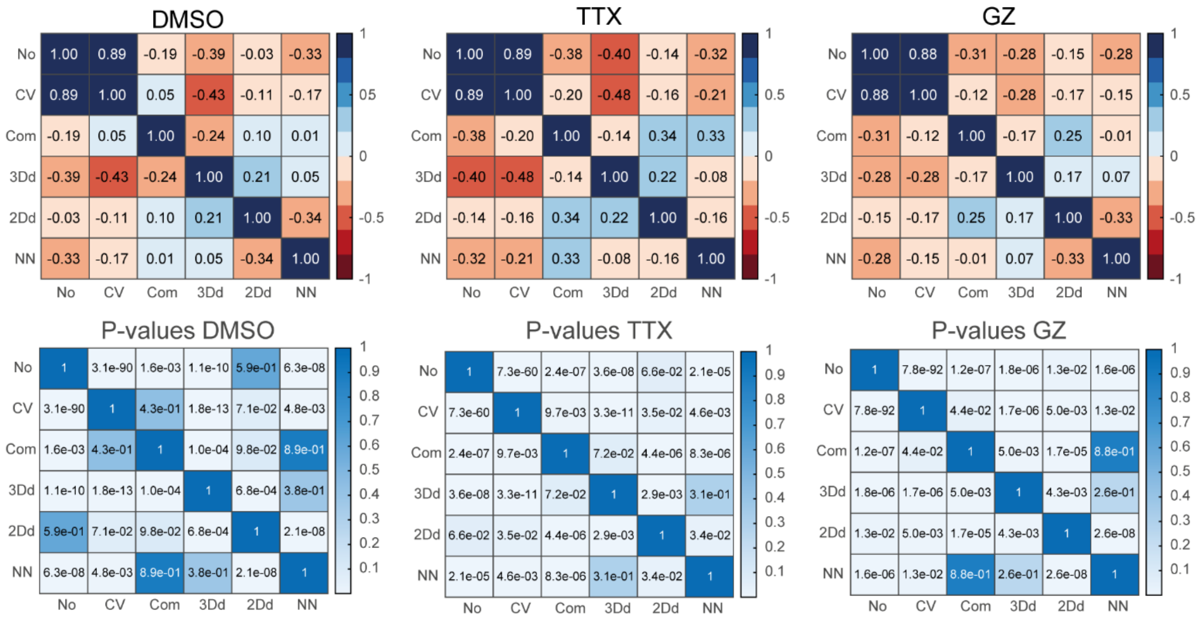


**Fig. S12. Correlation matrices for treated samples and corresponding p-values.**

Upper raw: correlation matrixes of DMSO, TTX and GZ treated samples. Color-coded values represent the Spearman correlation coefficients.

Lower raw: p-values to assess the goodness of the correlation.

Same dataset as in Fig. 4.

**Table S1. Performances of different labeling strategies.**

Quantification of the performances of samples labeled with either the nanobody conjugated to SulfoCy5 or to DNA-PAINT, or a primary antibody and secondary nanobody for DNA-PAINT. DBSCAN performed with eps = 200 nm and *minPoints* = 40. TIDs stands for Trace IDs, IQR for the interquartile range, n for the number of synaptic sites identified and N for the total number of images analyzed.

| **Parameter** | **MEDIAN** | **IQR** | **MEDIAN** | **IQR** | **MEDIAN** | **IQR** |
|---|---|---|---|---|---|---|
| | **Nanobody SulfoCy5** (n = 42, N =6) | | **Nanobody DNA-PAINT** (n = 167, N = 8) | | **Antibody + 2ry nanobody DNA-PAINT** | |
| **Avg. number synaptic sites / image** | 7 | | 21 | | 27 | |
| **TIDs / synaptic site** | 86.5 | 156 | 268 | 205.75 | 181 | 191 |
| **Units / synaptic site** | 57 | 108 | 117 | 93.5 | 90 | 91.75 |

**Table S2. Correlation coefficients and corresponding p-values of the matrix shown in Fig. 3g**

| Coefficients | | | | | | | | P-values | | | | | | |
|---|---|---|---|---|---|---|---|---|---|---|---|---|---|---|
| | No | CV | Comp | 3Dd | 2Dd | NN | | | No | CV | Comp | 3Dd | 2Dd | NN |
| **No** | 1.000 | 0.853 | -0.263 | -0.370 | -0.125 | -0.397 | | **No** | 1.000 | 0.000 | 0.000 | 0.000 | 0.037 | 0.000 |
| **CV** | 0.853 | 1.000 | 0.002 | -0.432 | -0.100 | -0.222 | | **CV** | 0.000 | 1.000 | 0.978 | 0.000 | 0.097 | 0.000 |
| **Comp** | -0.263 | 0.002 | 1.000 | -0.237 | 0.114 | 0.113 | | **Comp** | 0.000 | 0.978 | 1.000 | 0.000 | 0.058 | 0.060 |
| **3Dd** | -0.370 | -0.432 | -0.237 | 1.000 | 0.368 | -0.046 | | **3Dd** | 0.000 | 0.000 | 0.000 | 1.000 | 0.000 | 0.445 |
| **2Dd** | -0.125 | -0.100 | 0.114 | 0.368 | 1.000 | -0.050 | | **2Dd** | 0.037 | 0.097 | 0.058 | 0.000 | 1.000 | 0.402 |
| **NN** | -0.397 | -0.222 | 0.113 | -0.046 | -0.050 | 1.000 | | **NN** | 0.000 | 0.000 | 0.060 | 0.445 | 0.402 | 1.000 |

**Table S3. Detailed Statistics information about the chosen Linear Mixed Effect Model.** Parameter of interest, Model formula, the estimates, 95 % confidence interval and p-values relative to the reference (DMSO). Round refers to the experimental round (*i.e.* experimental replicate). File_ID refers to the image ID. The number of analyzed PSD clusters were (n = number of PSD-95 clusters, N = independent experimental round): DMSO: n = 261, N = 4; TTX: n = 174, N = 4; GZ: n = 279, N = 3.

| Parameter | Model Formula | Treatment | Estimate | 95% CI [Lower - Upper] | p-value |
|---|---|---|---|---|---|
| Cluster Volume (CV) | CV ~ Treatment +(1\|Round:File_ID) | DMSO | 5.71E-22 | [3.25E-22 - 8.18E-22] | |
| | | GZ | 5.76E-22 | [2.30E-22 - 9.23E-22] | 0.98 |
| | | TTX | 6.92E-22 | [3.03E-22 - 1.08E-21] | 0.54 |
| Projected Area (A) | A ~ Treatment +(1\|Round:File_ID) | DMSO | 2.55E-14 | [1.65E-14 - 3.46E-14] | |
| | | GZ | 2.66E-14 | [1.38E-14 - 3.93E-14] | 0.87 |
| | | TTX | 3.69E-14 | [2.26E-14 - 5.12E-14] | 0.12 |
| Units per Cluster (UC) | UC ~ Treatment +(1\|Round:File_ID) | DMSO | 79.1 | [5.40E+01 - 1.04E+02] | |
| | | GZ | 75.96 | [4.06E+01 - 1.11E+02] | 0.86 |
| | | TTX | 94.12 | [5.46E+01 - 1.34E+02] | 0.46 |
| Units 3D density (3Dd) | 3Dd ~ Treatment +(1\|Round:File_ID) | DMSO | 2.69E+23 | [2.36E+23 - 3.02E+23] | |
| | | GZ | 2.59E+23 | [2.13E+23 - 3.06E+23] | 0.67941 |
| | | TTX | 2.47E+23 | [1.94E+23 - 3.00E+23] | 0.40808 |
| Units 2D density (2Dd) | 2Dd ~ Treatment +(1\|Round:File_ID) | DMSO | 3.43E+15 | [3.18E+15 - 3.85E+15] | |
| | | GZ | 3.45E+15 | [3.19E+15 - 3.72E+15] | 0.84 |
| | | TTX | 3.01E+15 | [2.71E+15 - 3.30E+15] | 0.01 |
| Compactness (Com) | Com ~ Treatment +(1\|Round:File_ID) | DMSO | 0.055 | [4.97E-02 - 5.96E-02] | |
| | | GZ | 0.053 | [4.56E-02 - 5.93E-02] | 0.53 |
| | | TTX | 0.044 | [3.66E-02 - 5.21E-02] | 0.01 |
| TIDs per Cluster (TidC) | TidC ~ Treatment +(1\|Round:File_ID) | DMSO | 186.27 | [1.14E+02 - 2.58E+02] | |
| | | GZ | 171.94 | [7.01E+01 - 2.74E+02] | 0.78 |
| | | TTX | 196.23 | [8.30E+01 - 3.09E+02] | 0.86 |
| TIDs 3D density (T3Dd) | T3Dd ~ Treatment +(1\|Round:File_ID) | DMSO | 6.29E+23 | [5.28E+23 - 7.30E+23] | |
| | | GZ | 6.01E+23 | [4.61E+23 - 7.42E+23] | 0.7 |
| | | TTX | 4.66E+23 | [3.06E+23 - 6.26E+23] | 0.046 |
| TIDs 2D density (T2Dd) | T2Dd ~ Treatment +(1\|Round:File_ID) | DMSO | 7.94E+15 | 7.18E+15 - 8.70E+15] | |
| | | GZ | 7.76E+15 | [6.71E+15 - 8.82E+15] | 0.75 |
| | | TTX | 6.28E+15 | [5.08E+15 - 7.48E+15] | 0.007 |
| Units Nearest Neighbour (UNN) | UNN ~ Treatment +(1\|Round:File_ID) | DMSO | 1.76E-08 | [1.70E-08 - 1.81E-08] | |
| | | GZ | 1.78E-08 | [1.70E-08 - 1.85E-08] | 0.55 |
| | | TTX | 1.80E-08 | [1.72E-08 - 1.88E-08] | 0.28 |
| TIDs Nearest Neighbour (TNN) | TNN ~ Treatment +(1\|Round)+(1\|Round:File_ID) | DMSO | 7.17E-09 | [6.51E-09 - 7.82E-09] | |
| | | GZ | 7.43E-09 | [6.57E-09 - 8.30E-09] | 0.54 |
| | | TTX | 8.15E-09 | [7.20E-09 - 9.10E-09] | 0.043 |

**Movie S1.**

Example of a 3D MINFLUX data. Confocal image of phalloidin (white) overlayed with the identified PSD-95 clusters (purple), zoom in to 4 selected clusters and corresponding 3D rendering. TIDs rendered as gaussian with FWHM of 12 nm. Size of the confocal field of view: 15,36 x 18,16 µm.
